# Spindle assembly checkpoint strength is governed by cell size and PAR-mediated cell fate determination in *C. elegans*

**DOI:** 10.1101/134809

**Authors:** Abigail R. Gerhold, Vincent Poupart, Jean-Claude Labbé, Paul S. Maddox

**Author notes:** Correspondence to: Abigail R. Gerhold.

## Abstract

The spindle assembly checkpoint (SAC) is a conserved mitotic regulator that preserves genome stability. Despite its central role in preserving the fidelity of mitosis, the strength of the SAC varies widely between cell types. How the SAC is adapted to different cellular contexts remains largely unknown. Here we show that both cell size and cell fate impact SAC strength. While smaller cells have a stronger SAC, cells with a germline fate show increased SAC activity relative to their somatic counterparts across all cell sizes. We find that enhanced SAC activity in the germline blastomere P_1_ requires proper specification of cell fate downstream of the conserved PAR polarity proteins, supporting a model in which checkpoint factors are distributed asymmetrically during early germ cell divisions. Our results indicate that size scaling of SAC activity is modulated by cell fate and reveal a novel interaction between asymmetric cell division and the SAC.

## Introduction

The fidelity of mitosis depends upon equal partitioning of the replicated genome between daughter cells. During mitosis, sister chromatid pairs connect to the mitotic spindle via kinetochore-microtubule attachments. Stable attachment of sister chromatids to opposite spindle poles (bi-orientation) ensures that, upon chromatid separation, one copy segregates to each daughter cell. Attachment of sister chromatids to the mitotic spindle is an inherently stochastic process of variable duration (1). Thus to safeguard against chromosome segregation errors, the spindle assembly checkpoint (SAC) monitors kinetochore-microtubule attachments and prevents anaphase onset until stable bi-orientation has been achieved. Weakening of the SAC can lead to aneuploidy and has been associated with tumour development in both model systems and human cancers (2, 3). Accordingly, a major class of anti-mitotic chemotherapeutics (spindle poisons) depends upon robust SAC activity to disrupt mitosis in cancer cells (4).

Interfering with the formation of stable kinetochore-microtubule attachments prevents cells from satisfying the SAC and leads to prolonged mitotic delays. However, the SAC cannot block anaphase indefinitely, and most cells, even in the complete absence of spindle microtubules, will eventually exit mitosis (5). The duration of mitotic arrest under conditions that preclude satisfaction of the SAC is often used as an indication of the “strength” of the SAC. The strength of the SAC varies widely between different cell types (6, 7), cells from different organisms (5) and cells at different developmental stages (8-10). While variation in the strength of the SAC is widespread, an understanding of how this variation is achieved is currently lacking.

The core SAC module consists of the proteins Bub1, Bub3, BubR1, Mad1 and Mad2, which are recruited to unattached kinetochores in a stepwise fashion to promote assembly of the mitotic checkpoint complex (MCC), which acts as a diffusible “wait-anaphase” signal (11, 12). The MCC consists of Mad2, BubR1 and Bub3 bound to the anaphase-promoting complex/cyclosome (APC/C) cofactor Cdc20 (13). The MCC binds to the APC/C (14, 15), blocking degradation of APC/C substrates, namely Cyclin B (16) and securin (17), thereby maintaining a mitotic state. Once all kinetochores are stably bound by microtubules, disassembly of the MCC (18, 19), dephosphorylation of SAC proteins (20) and stripping of Mad1 and Mad2 from all kinetochores (21) effectively silences the SAC and permits anaphase onset and mitotic exit.

Mitotic exit, despite failure to satisfy the SAC, occurs when activity of Cyclin B/Cdk1 falls below the threshold necessary to maintain a mitotic state (5). In mammalian cells, this mitotic “slippage” occurs because the active cytoplasmic pool of MCC is not sufficient to completely inhibit the APC/C, and the progressive degradation of Cyclin B eventually enables mitotic exit (22). The steady-state concentration of the MCC pool is influenced by the rate of new, kinetochore-catalyzed MCC generation and disassembly of existing cytoplasmic MCC (23, 24), the former being intimately related to the severity of the spindle defect (25-27). Thus variation in the strength of the SAC, as assayed by the duration of mitotic delay following spindle perturbation, may be linked to differences in MCC production, activity or stability.

One factor that contributes to SAC strength in certain circumstances is cell size. *In vitro* experiments using *Xenopus laevis* egg extracts suggested that an increased nuclear to cytoplasmic ratio, as would be found in smaller cells, could increase SAC activity (28). Recent work in *C. elegans* embryos has shown that the strength of the SAC indeed scales with cell size, with smaller cells exhibiting longer SAC-dependent delays upon spindle perturbation (29). However, in other organisms, the SAC remains inactive until the mid-blastula transition and acquisition of SAC activity is neither accelerated by decreasing cell volume (*Xenopus laevis*, (8, 9)) nor delayed by increasing cell volume (*Danio rerio*, (10)), indicating that SAC activity can also be developmentally regulated independently of changes in cell volume. How this developmental regulation of the SAC is achieved is unknown.

We have recently reported that the adult germline stem cells (GSCs) of *C. elegans* exhibit a stronger SAC relative to early embryonic cells (30) providing a tractable model in which to examine SAC adaptability. Here we use an inducible monopolar spindle assay to investigate the developmental origins of enhanced germline SAC activity. In agreement with Galli and Morgan (29), we find that the duration of SAC-dependent mitotic delays increases as cell size decreases during embryogenesis. However, the relationship between cell size and SAC activity is strongly influenced by cell fate, with cells in the germline lineage displaying a stronger SAC relative to their cell size than cells with a somatic fate. At the 2-cell stage, we find that differential SAC activity in the somatic AB versus germline P_1_ blastomere requires the highly conserved PAR polarity proteins, supporting a model in which asymmetric partitioning of a checkpoint-enhancing factor increases the strength of the SAC in germline cells.

## Results

### Germline blastomeres are more sensitive to spindle perturbations

*C. elegans* GSCs are derived from a single founder cell (P_4_), which is specified during embryogenesis by a series of asymmetric cell divisions (31). As the *C. elegans* embryonic lineage is invariant and fully mapped, each cell can be identified by its position within the embryo and its cell cycle characteristics (Figure 1A, (31, 32)). We were therefore able to ask whether the embryonic precursors to GSCs, cells in the germline P lineage, also exhibited an enhanced SAC, by comparing the duration of mitosis between each of the founding cell lineages, in cells with or without spindle perturbations. As *C. elegans* embryos are largely refractory to treatment with small molecule spindle poisons without physical or genetic manipulations to permeabilize the egg shell (33, 34), we opted for a genetic method to induce spindle perturbations with temporal control. We utilized temperature sensitive alleles of genes involved in spindle formation, combined with fluorescent markers to visualize the mitotic spindle (β-tubulin fused to GFP, hereafter β-tubulin::GFP) and/or the nucleus/chromatin (histone H2B fused to mCherry, hereafter H2B::mCH). This allowed us to disrupt spindle formation with a simple temperature shift using a standard temperature-controlled imaging chamber and to follow mitotic progression by monitoring changes in nuclear and chromosome morphology.

**Figure 1.**
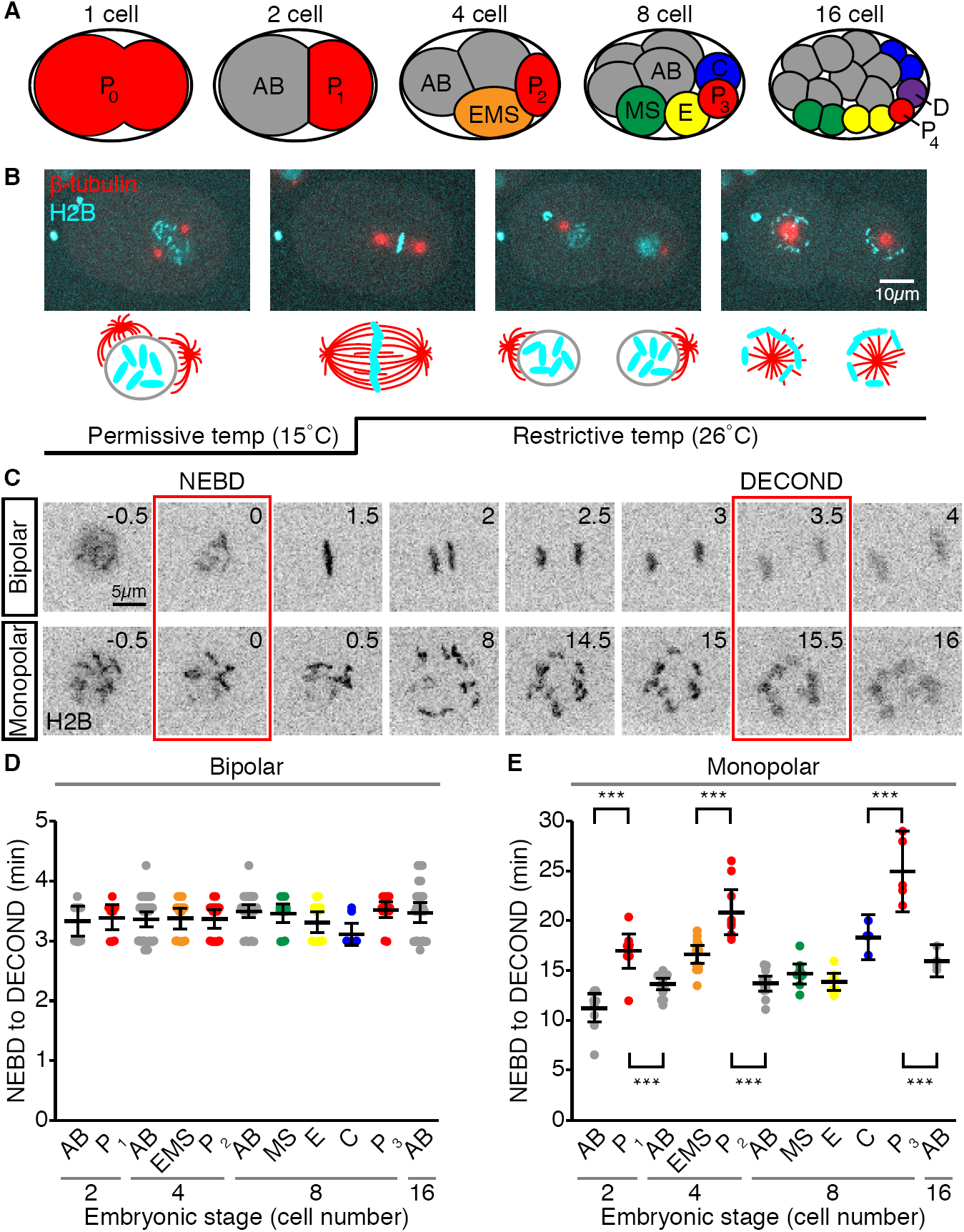
Germline blastomeres have longer mitotic delays than somatic cells. (A) Schematic depiction of the first four rounds of division in *C. elegans* embryogenesis, with the germline (P) lineage in red. (B) Representative images of a 1- to 2-cell stage *zyg-1(or297ts)* embryo expressing H2B::mCH and β-tubulin::GFP, shifted from permissive (15°C) to restrictive (26°C) temperature, showing a normal bipolar division at the 1-cell stage (P_0_) and monopolar spindles at the 2-cell stage (AB and P_1_). Embryo is oriented with the anterior pole to the left. (C) Single cropped time-lapse images showing a bipolar (top) and monopolar (bottom) mitosis in two P_1_ blastomeres, from nuclear envelop breakdown (NEBD) when non-chromatin associated H2B::mCH is lost, to chromosome decondensation (DECOND) when bright, compact H2B::mCH-marked chromatin is lost. (D and E) Graphs showing the duration of bipolar (D) and monopolar (E) mitoses in different embryonic cells from 2 to 16-cell stage embryos. Black bars represent the mean. Error bars show the 95% confidence interval for the mean. Color-coding of other cells is as in (A). Germline blastomeres (red) with monopolar spindles remain in mitosis for significantly longer than somatic cells at the same and at later embryonic stages (*** = p < 0.001 by an Anova1 with Tukey-Kramer post-hoc test).

We used a recessive temperature-sensitive allele of the gene encoding the polo-related kinase ZYG-1, which prevents centriole duplication and produces cells with monopolar spindles (*zyg-1(or297)*, hereafter *zyg-1(ts*), (35, 36)). Monopolar spindles can generate mitotic delays that are indistinguishable from Nocodazole treatments that either partially (6, 7) or completely (25) inhibit microtubule formation, suggesting a robust SAC response, and are an established means for SAC studies in *C. elegans* (37). *zyg-1(ts)* is fast-acting (36) and readily permitted the induction of monopolar spindles in all embryonic cells up to the 16-cell stage. Following NEBD, condensed chromosomes spread radially around the single spindle pole and remained dispersed until the start of decondensation (Figures 1B and 1C, Movie S1). In later stage embryos, centrosome duplication was unaffected by temperature shift and all divisions were bipolar (data not shown).

In agreement with previous reports, we found that the duration of mitosis in cells without spindle perturbations was invariant across embryonic stages and cell lineages (Figure 1D, (29, 38)). In cells in which spindle formation was perturbed, we noticed significant differences in the duration of mitosis between cells at different embryonic stages and in different lineages. Notably, cells in the germline P lineage remained in mitosis for significantly longer than both their immediate somatic siblings and somatic cells at later embryonic stages (Figures 1E). A similar result was obtained using a temperature sensitive allele of the microtubule subunit β-tubulin (*tbb-2(or362*), (39)), which alters microtubule dynamics and/or stability, to disrupt spindle formation (Figure S1 and Movie S2)

### The difference in mitotic delay between germline and somatic cells is not solely due to cell size

Specification of the germline is achieved via a series of asymmetric cell divisions, such that, the germline blastomere is always smaller than its immediate somatic sibling (31). As the strength of the SAC negatively correlates with cell size during *C. elegans* embryogenesis (29), the increased duration of monopolar mitoses in germline blastomeres could be due to their relatively small size. Thus, we considered cell size in two ways. We used nuclear area, which scales with cell size in many organisms including *C. elegans* (40, 41), and which we could measure for the same cells in which we assayed mitotic delay, as a proxy for cell size (Figure S2A and S2B). In both the germline P and somatic AB lineages, the duration of monopolar mitoses negatively correlated with nuclear area, supporting the idea that smaller cells possess a stronger SAC (r = -0.73, p < 0.001 and r = -0.61, p < 0.001, respectively). However, germline cells exhibited longer mitotic delays than somatic AB lineage cells with similar nuclear areas (Figure 2A), suggesting that, between comparably sized cells, the SAC is stronger in germline cells.

**Figure 2.**
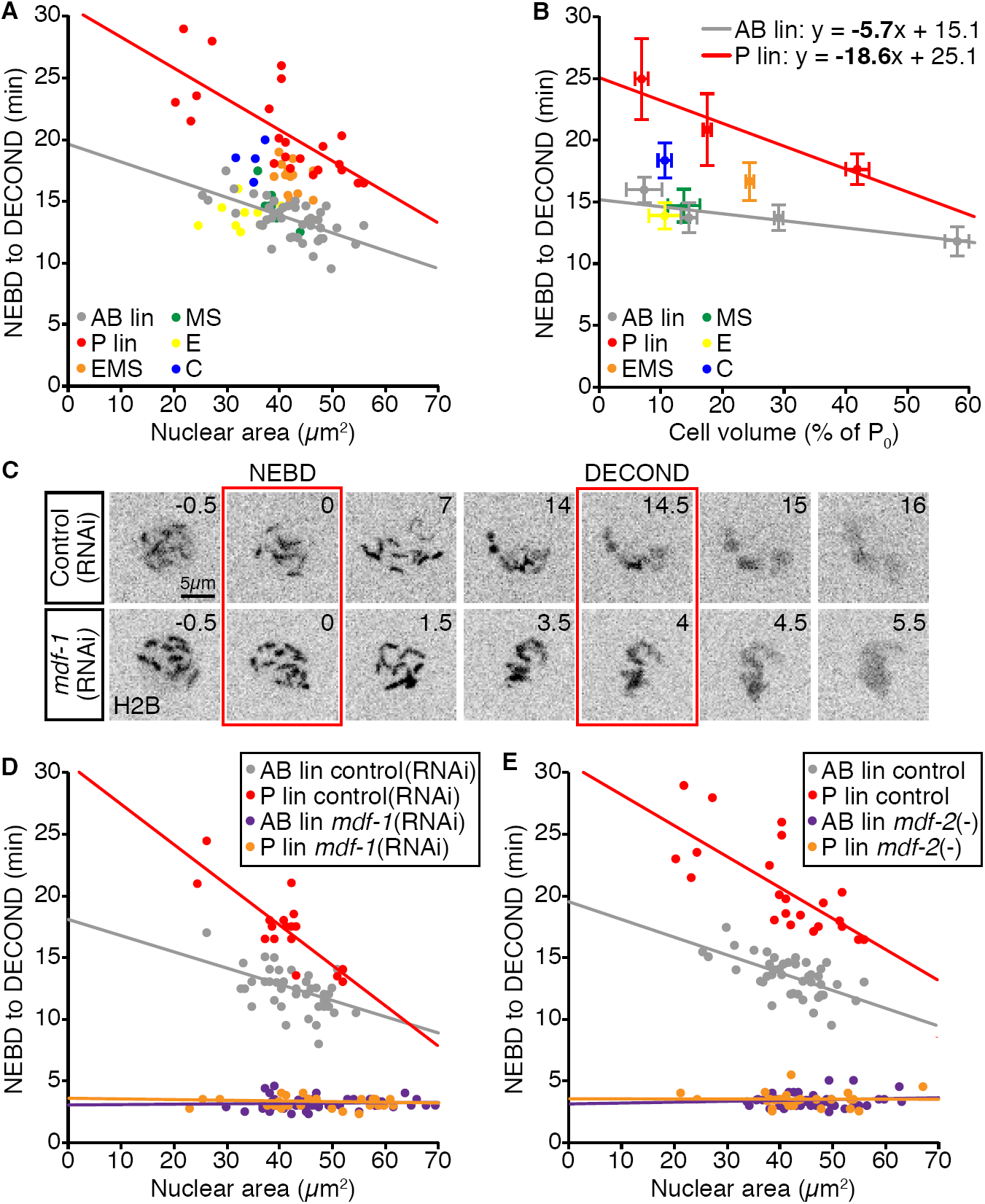
The duration of monopolar mitoses in germline blastomeres are disproportionate to their cell size and require the SAC. (A) Scatter plot showing the relationship between nuclear area (as a proxy for cell size; see Figure S2) and the duration of monopolar mitoses in different embryonic cells. Lines show the linear least squares regression fit for germline P lineage cells (red, r = -0.73, p < 0.001) versus somatic AB lineage cells (grey, r = -0.61, p < 0.001). (B) Graph showing the relationship between mean cell volume (calculated separately as a proportion of starting embryonic (P_0_) volume) and mean duration of monopolar mitoses for the cells shown in (A). Error bars show standard deviation. Linear least squares regression models for germline P (red, r = -0.72, p < 0.001) and somatic AB (grey, r = -0.62, p < 0.001) lineages were calculated by plotting the duration of mitosis in each cell by the average volume for all cells of its respective lineage/cell stage. Statistically different regression coefficients are written in bold and were compared using a non-parametric bootstrap (p = 0.02 for slope). For (A) and (B), p-values for Pearson’s coefficient (r) were determined by Student’s t-test. (C) Single cropped time-lapse images showing nuclear envelop breakdown (NEBD) and chromosome decondensation (DECOND) in monopolar germline P_1_ cells from a Control(RNAi) (top) and a *mdf-1*(RNAi) (bottom) embryo. (D and E) Scatter plots showing the relationship between nuclear area and the length of monopolar mitoses in somatic AB and germline P lineage cells in control embryos (red = P/germline, grey = AB/somatic) compared to embryos in which SAC activity has been impaired (orange = P/germline, purple = AB/somatic). (D) *mdf-1*(RNAi) versus Control(RNAi). (E) *mdf-2*(null) versus *mdf-2*(+). Control data in (E) are reproduced from (A).

Nuclear size reached an upper limit and did not reflect differences in cell volume in larger cells (Figure S2B and S2F). A similar phenomenon has been described for the size scaling of both individual chromosomes and the mitotic spindle (42, 43). Thus, in a separate experiment, we calculated the average volume for each cell in the different lineages relative to the volume of the founding P_0_ blastomere (Figure S2C-E). In both lineages, mitotic delay increased as cell volume decreased (Figure 2B, r = -0.62, p < 0.001 for the AB lineage; r = -0.72, p < 0.001 for the P lineage). However, the relationship between cell volume and the duration of monopolar mitoses differed significantly between the two lineages (AB versus P regression slope: p = 0.03). Here again, germline cells experienced longer mitotic delays relative to their cell volume than somatic AB cells, suggesting that SAC strength is also shaped by cell lineage.

### Both cell size and lineage-specific differences in the duration of monopolar mitoses are checkpoint dependent

The increased duration of monopolar mitoses in germline blastomeres could be due to a stronger SAC; alternatively, checkpoint-independent factors could contribute to a delayed mitotic exit in these cells. To discriminate between these possibilities, we examined the duration of monopolar mitoses in the germline P and somatic AB lineages when checkpoint activity was eliminated using RNAi depletion of the *C. elegans* ortholog of Mad1, *mdf-1*(RNAi) (Figure 2C and 2D, (44)), or a null allele of the *C. elegans* ortholog of Mad2, *mdf-2(tm2190)* (Figure 2E, (45, 46)). In the absence of checkpoint activity, cells with monopolar spindles rapidly exited mitosis, with chromosome decondensation evident within 4 minutes of NEBD (Figure 2C and Movie S3). Plotting the duration of monopolar mitoses in the germline P and somatic AB lineages relative to cell size (as approximated by nuclear area; see Figure S2 and preceding section) revealed that all cells, irrespective of their size or lineage, exited mitosis with the same timing (Figure 2D and 2E). The duration of checkpoint deficient monopolar mitoses was comparable to that of normal bipolar divisions (3.3±0.5 versus 3.4±0.3 minutes, respectively). Thus monopolar spindle-induced mitotic delays are entirely checkpoint dependent and differences in the duration of these delays are likely to reflect differences in the strength of the SAC.

### SAC activity may also be subject to developmental regulation

The strength of the checkpoint increases in both germline and somatic AB cells as cell size decreases. If cell size affected SAC strength according to a universal size-scaling relationship, the linear regression lines for mitotic delay versus cell volume for each lineage should be parallel. Instead, SAC activity increases more rapidly with decreasing cell volume in the germline P lineage (Figure 2B and S3A), suggesting that either cell size impacts SAC activity differently in the two lineages or that SAC activity is progressively modified during lineage development. To distinguish between these possibilities, we modified embryo size experimentally and measured the duration of monopolar mitoses at a single developmental stage, in the somatic AB and germline P_1_ blastomeres of 2-cell stage embryos.

We used RNAi depletion of ANI-2 and PTC-1 to generate small embryos, and C27D9.1 to generate large embryos (Figure 3A and data not shown, (41, 47-49)). Altering embryo size by these means gave us a 5-fold range of embryo volumes, from 1.2 × 10^4^ μm^3^, slightly smaller than a control P_1_ cell, to 6.1 × 10^4^ μm^3^, about twice the size of an average control embryo, without disrupting basal mitotic timing and other defining features of AB and P_1_, such as cell cycle asynchrony and cell size asymmetry (Figure S3B-D). Hereafter cell volume is expressed as the radius of the equivalent sphere (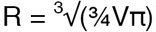), which we will call “cell size”.

**Figure 3.**
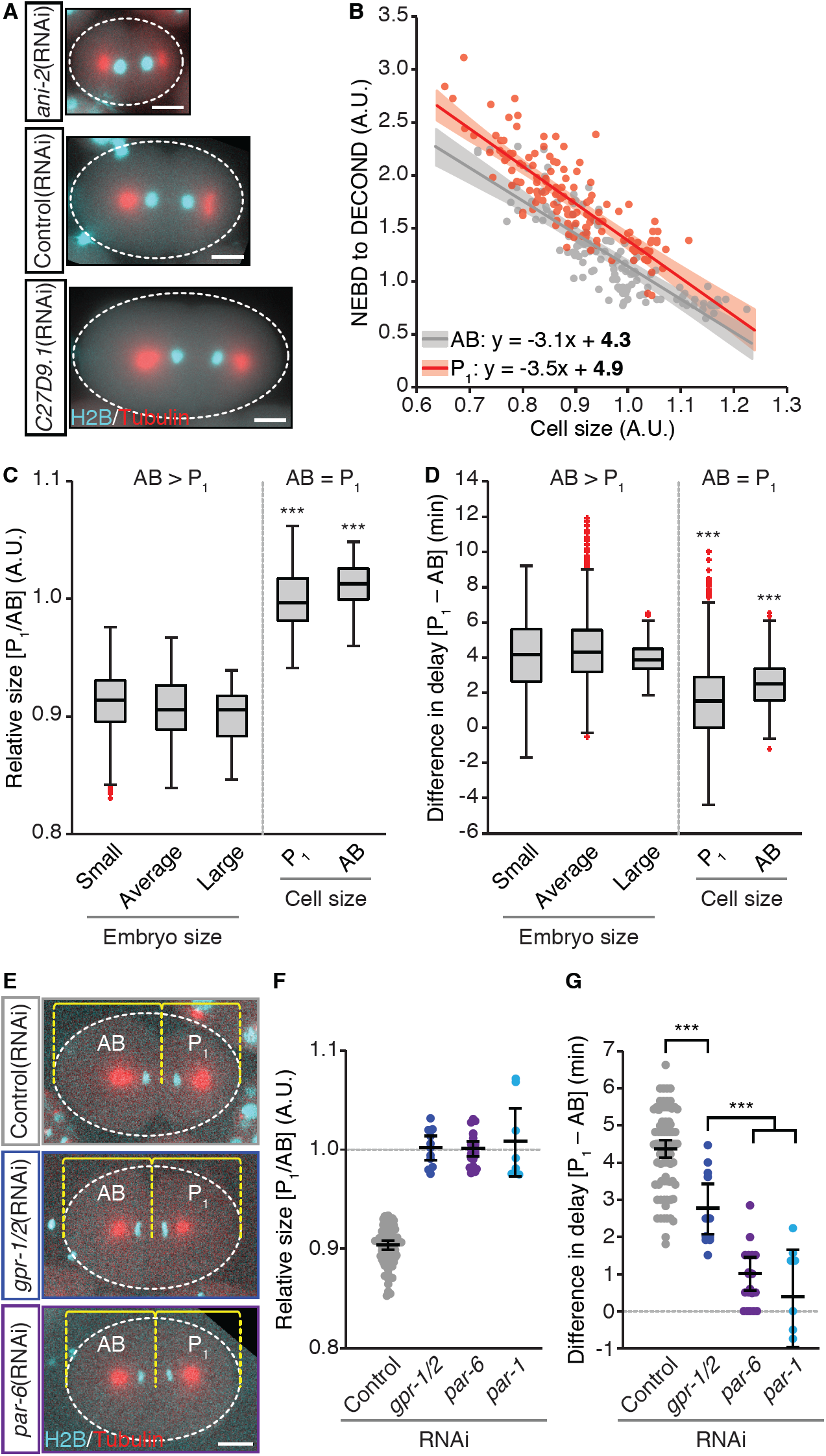
Longer monopolar mitoses in P_1_ germline blastomeres depend on cell size and cell fate. (A) Representative images of *zyg-1(or297ts),* H2B::mCH, β-tubulin::GFP embryos following RNAi depletion of ANI-2, which gives small embryos, or C27D9.1, which gives large embryos, as compared to control. (B) Scatter plot showing the duration of monopolar mitoses relative to cell size in somatic AB (grey) and germline P_1_ (red) cells following RNAi-induced changes in embryo volume. Cell volumes were converted into the radius of the corresponding sphere (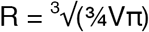), which we call “cell size”. Both timing of NEBD to DECOND and cell size were normalized to the mean value for Control(RNAi) AB cells. Lines represent the linear least squares regression fit with 95% confidence interval (shaded regions). Linear regression models are shown with statistically different coefficients in bold. Regression coefficients were compared using a non-parametric bootstrap (p = 0.12 for slope and 0.008 for y-intercept). (C and D) Comparison of the duration of monopolar mitoses in somatic AB and germline P_1_ cells with a normal cell size ratio (AB > P_1_), in small, average and large embryos, versus AB and P_1_ cells that are both the same size (AB = P_1_) as Control(RNAi) P_1_ cells (“P_1_”) or Control(RNAi) AB cells (“AB”). Box plots represent the outcome for all possible comparisons within a class. Box edges are the 25th and 75th percentiles, whiskers extend to the most extreme data points. Outliers are plotted individually in red. Asterisks indicate statistical significance relative to all AB > P_1_ classes. (C) Cell size of P_1_ relative to AB. (D) Difference in the duration of monopolar mitoses between AB and P_1_. (E) Representative images of *zyg-1(or297ts)*, H2B::mCH, β-tubulin::GFP embryos treated with Control(RNAi), *gpr-2*(RNAi) or *par-6*(RNAi), showing the position of the spindle midpoint along the anterior-to-posterior axis of P_0_. (F-G) Graphs showing the relative size (F) and difference in the duration of monopolar mitoses (G) for AB/P_1_ sibling pairs following RNAi depletion of GPR-2 (blue), PAR-6 (purple), and PAR-1 (light blue). Control embryos are shown in grey. Black bars represent the mean. Error bars show the 95% confidence interval for the mean. For (C), (D), (F) and (G) *** = p < 0.001 by an Anova1 with Tukey-Kramer post-hoc test. For (A) and (E) Scale bar is 10μm. Anterior is to the left.

Cell size and the duration of monopolar mitoses were negatively correlated in both AB and P_1_ blastomeres (Figure 3B, r = -0.86, p < 0.001 and r = -0.83, p < 0.001, respectively), indicating that cell size can impact SAC activity irrespective of developmental stage. While the rate at which the duration of monopolar mitoses changed relative to cell size was similar for AB and P_1_ (Figure 3B, AB versus P_1_ regression slope: p = 0.12), the average duration of monopolar mitoses was always higher in P_1_ cells (Figure 3B, AB versus P_1_ regression y-intercept: p = 0.008), indicating that P_1_ cells have a stronger SAC regardless of cell size. These results suggest that cell identity sets a baseline of SAC activity, which is then scaled according to cell size similarly in AB and P_1_ and argue against a cell fate-specific difference in SAC size scaling during development of the AB and P lineages. Thus we favor the alternative hypothesis, that checkpoint activity is modified within each lineage as development progresses. Altogether our results demonstrate that SAC activity is determined by a combination of cell size, cell fate and potentially developmental stage in the early *C. elegans* embryo.

### PAR protein-mediated cytoplasmic asymmetries are required for differential SAC activity in AB versus P_1_

The pattern of checkpoint activity that we have observed in 2- to 16-cell stage embryos mirrors the asymmetric inheritance of certain germline factors, raising the possibility that a determinant of SAC activity could be similarly regulated. For example, several germline proteins are initially present at low levels in the somatic siblings of germline blastomeres, but absent in their progeny (50-53), and germline-enriched maternal transcripts are initially symmetric during cell division and subsequently lost from somatic cells and/or their progeny (54). Correspondingly, the sibling of the germline blastomere in 4- and 8-cell stage embryos (EMS and C, respectively) exhibited monopolar mitoses that were intermediate in duration to those in germline P and somatic AB lineage cells, and, in the progeny of EMS (MS and E), these longer monopolar mitoses were lost (Figures 1E and 2B). The asymmetric inheritance of germline determinants is regulated by the highly conserved PAR proteins, which lead us to ask whether differences in SAC activity also require PAR proteins.

The polarized, cortical distribution of PAR proteins regulates both cell size and the cytoplasmic distribution of cell fate determinants during the division of P_0_ (55). PAR-6, PAR-3 and the atypical protein kinase C PKC-3 localize to the anterior cortex, while PAR-2 localizes to the posterior cortex. The asymmetric cortical distribution of PAR proteins depends on mutual antagonism between the anterior and posterior PARs. Consequently, in the absence of anterior PARs, posterior PARs move into the anterior, and vice versa (55). The PAR-1 kinase is also enriched in the posterior and is both cytoplasmic and cortical (56). Activity of PAR-1 regulates the asymmetric segregation and/or degradation of cytoplasmic cell fate determinants, leading to the enrichment of germline factors in P_1_ and somatic factors in AB (57). PAR-1 plays a minimal role in the regulation of cortical PAR protein asymmetries (58-60), but is absolutely required for downstream cytoplasmic asymmetries (56, 61, 62).

To determine how much of the difference in SAC strength between AB and P_1_ was due to cell size, we compared the average difference in mitotic delay between AB and P_1_ cells that were the same size versus AB and P_1_ cells where the normal cell size ratio was preserved. Monopolar mitoses were 4.3±1.8 minutes longer, on average, in P_1_ cells when AB was larger than P_1_, irrespective of embryo volume, and 2.0±1.9 minutes longer, on average, in P_1_ when AB and P_1_ were the same size (Figure 3C and 3D). When we compared sibling AB/P_1_ pairs from control embryos versus embryos in which AB and P_1_ were the same size but maintained somatic versus germline fate (*gpr-1/2(RNAi*) (63-65)), the difference in the duration of monopolar mitoses was reduced from 4.4±1.1 to 2.7±1.0 minutes (Figure 3E and 3F and Movie S4). Thus, in 2-cell stage embryos, approximately half of the difference in checkpoint strength between AB and P_1_ is due to their size difference.

We next asked whether the activity of PAR proteins was responsible for the remaining difference in mitotic delay between equally sized AB and P_1_ cells. While disrupting PAR proteins in P_0_ will change the fate of the resulting cells, for simplicity we will continue to refer to the “anterior” blastomere as AB and the “posterior” blastomere as P_1_. RNAi depletion of PAR-6 resulted in sibling AB and P_1_ cells that were the same size, similar to *gpr-1/2*(RNAi) embryos (Figure 3E and 3F). However, in *par-6*(RNAi) embryos the difference in duration of monopolar mitoses was further reduced to 1.0±0.9 minutes, with several AB/P_1_ sibling pairs exiting mitosis synchronously (Figure 3G, Movie S5). Similarly, RNAi depletion of PAR-1 reduced the difference in the duration of monopolar mitoses between AB/P_1_ sibling pairs to 0.34±1.6 minutes (Figure 3G). RNAi depletion of additional PAR proteins (PKC-3 and PAR-2) gave similar results (Figure S4A and S4B). Thus PAR protein mediated asymmetries account for much of the remaining difference in checkpoint activity when AB and P_1_ are the same size. Depletion of PAR-6 and PAR-1 diminished the difference in mitotic delay between AB and P_1_ sibling pairs to a similar extent. As asymmetries in the cortical localization of PAR proteins are largely maintained in the absence of PAR-1 (58-60), while cytoplasmic asymmetries are lost entirely (56, 61, 62), our results suggest that PAR-1-dependent cytoplasmic asymmetries, but not cortical polarity per se, drive differential checkpoint activity in AB and P_1_.

### Loss of either anterior or posterior PAR proteins increases SAC activity in AB

PAR proteins regulate the levels and localization of asymmetrically distributed cellular components largely via the posterior enrichment of PAR-1. Removing anterior PAR proteins permits PAR-1 activity in the anterior, resulting in two cells with germline P_1_ traits. Conversely, eliminating posterior PAR proteins allows somatic AB traits to emerge in both cells (55). To ask whether SAC activity was subject to similar regulation, we measured the absolute duration of monopolar mitoses in PAR protein-depleted AB and P_1_ cells. The mean duration of monopolar mitoses relative to mean cell size for *par*(RNAi) AB and P_1_ cells (excepting *par-2*(RNAi)), more closely resembled the predicted values for P_1_ than AB control cells; whereas values for *gpr-1/2*(RNAi) embryos, in which polarity is normal, were similar to the corresponding control cell (Figure 4A). RNAi depletion of PAR proteins impacts the size of AB and P_1_, generating AB cells that are slightly smaller and P_1_ cells that are slightly larger than control AB and P_1_ cells, respectively. Thus, to confirm this trend, we utilized data from Figure 3B (hereafter referred to as “control” cells), binned cells by size and compared the duration of monopolar mitoses in *par*(RNAi) AB and P_1_ cells to comparably-sized, but normally polarized and fated, control AB and P_1_ cells.

**Figure 4.**
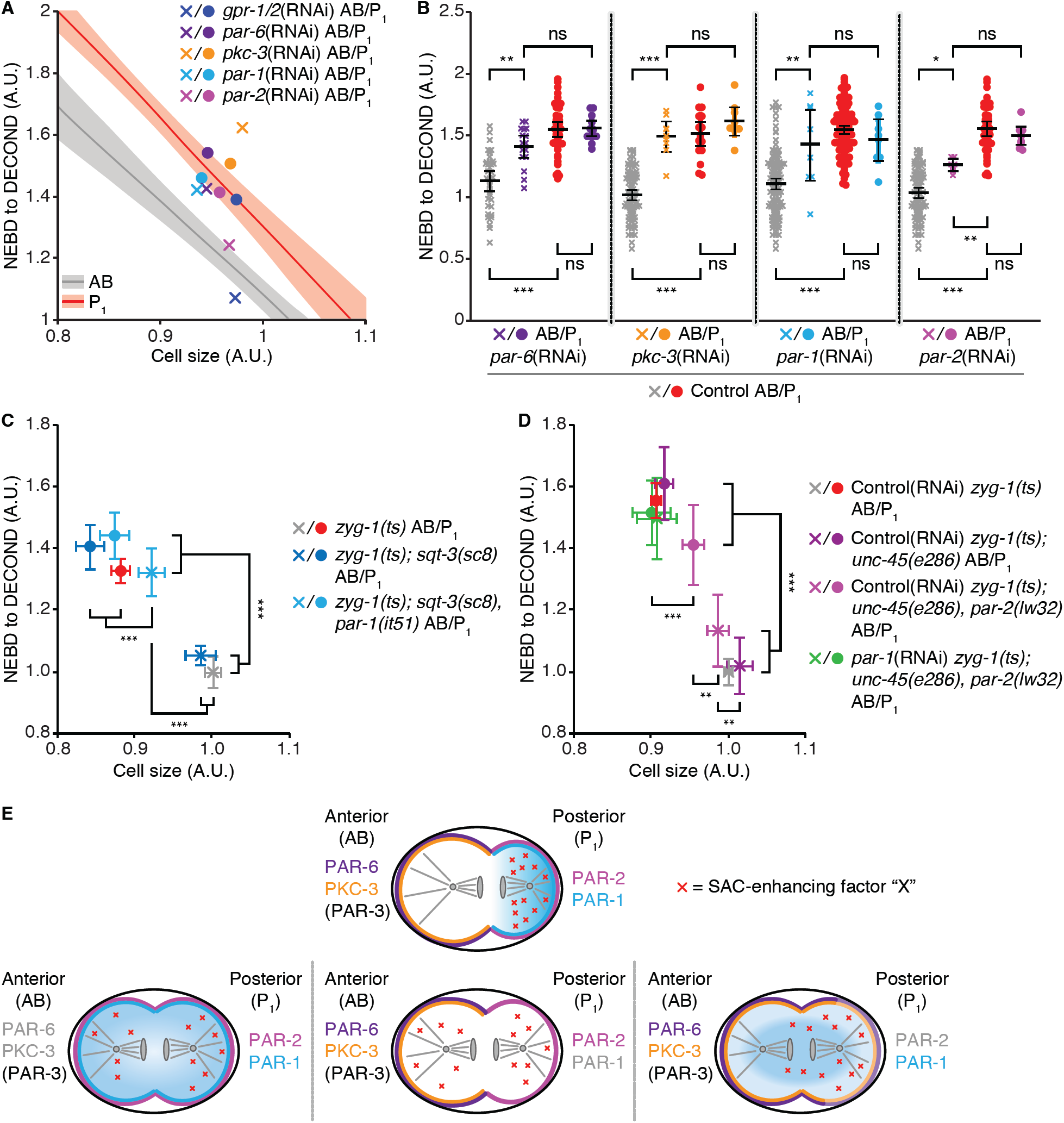
Depleting anterior PARs or PAR-1 increases the duration of monopolar mitoses in the somatic AB cell. (A) Graph plotting the mean duration of monopolar mitoses versus mean cell size for AB and P_1_ cells following RNAi depletion of GPR-2 (blue), PAR-6 (purple), PKC-3 (orange), PAR-1 (light blue) and PAR-2 (magenta). Lines represent the regression model with 95% confidence interval (shaded regions) for control somatic AB (grey) and germline P_1_ (red) cells and were generated from data in Figure 3D with additional data points from Control(RNAi) embryos. (B) Graph comparing the duration of monopolar mitoses in AB and P_1_ cells from PAR-depleted embryos with control somatic AB and germline P_1_ cells within the same size range. All PAR-depleted and control cells within ± 1 standard deviation (± 2 for *par-1*) of the mean size for all cells (AB and P_1_) for a given RNAi treatment are shown. Black bars represent the mean. Error bars show the 95% confidence interval for the mean. (C) Duration of monopolar mitoses versus cell size in control (grey and red), *sqt-3(sc8)* (dark blue), and *sqt-3(sc8) par-1(it51)* (light blue) AB and P_1_ cells. (D) Duration of monopolar mitoses versus cell size in control (grey and red), *unc-45(e286)* (purple)*, unc-45(e286) par-2(lw32)* (magenta) and PAR-1-depleted *unc-45(e286) par-2(lw32)* (green) AB and P_1_ cells. In (C) and (D) vertical brackets compare delay and horizontal brackets compare size. For (A-D) Xs = AB, circles = P_1_, cell size was derived from cell volume measurements as in Figure 3B. NEBD to DECOND and size measurements were normalized to the mean value for Control(RNAi) AB cells. p values were determined by an Anova1 with Tukey-Kramer post-hoc test, *** = p < 0.001, ** = p <0.01, * = p < 0.05 and ns = p > 0.05. (E) A model illustrating how localization of a checkpoint promoting factor (X) may be impacted by disrupting different PAR proteins. See main text for details.

RNAi depletion of the anterior PAR-6 and PKC-3 proteins increased the duration of monopolar mitoses in AB cells, such that mitotic delays in depleted AB cells closely resembled those in control P_1_ cells (Figure 4B). Surprisingly, however, RNAi depletion of the posterior PAR-1 protein also increased the overall duration of monopolar mitoses in AB, while depletion of the posterior PAR-2 protein gave an intermediate phenotype (Figure 4B). AB and P_1_ cells from *gpr-1/2*(RNAi) embryos, showed mitotic delays that were indistinguishable from comparably sized control AB and P_1_ cells, respectively (Figure S4C). All *par*(RNAi) embryos displayed the expected phenotypes with respect to cell cycle duration and asynchrony (66) (Figure S5A) indicating that our depletions were efficient.

We confirmed these results using a kinase-dead allele of *par-1* (*it51* (56)) and a near-null allele of *par-2* (*lw32* (58, 67)), with their respective genetic controls (see Table S1 for complete genotypes). AB cells in *par-1(it51)* embryos had monopolar mitoses similar in duration to control P_1_ cells and significantly longer than control AB cells (Figure 4C). In *par-2(lw32)* AB and P_1_ cells, the mean duration of monopolar mitoses was largely unchanged. RNAi depletion of PAR-1 in *par-2(lw32)* embryos eliminated the difference in mitotic delay between sibling AB/P_1_ pairs almost entirely (0.06 ± 0.9 minutes), and phenocopied *par-1(it51)* and *par-1*(RNAi) alone, with unaffected P_1_ cells and longer monopolar mitoses in AB cells (Figure 4CD). *par-1(it51)*, *par-2(lw32)* and *par-2(lw32); par-1*(RNAi) embryos all exhibited changes in cell cycle duration and asynchrony consistent with their genotype (Figure S5B). Under all conditions, *par-1(-)* AB cells were significantly smaller than control AB cells and comparable in size to control P_1_ cells (Figure 4C and 4D); however, in light of our AB/P_1_ size scaling and *gpr-1/2*(RNAi) results (Figure 3), small AB cells should have shorter monopolar mitoses than comparably sized control P_1_ cells. As monopolar mitoses in *par-1(-)* AB cells are largely indistinguishable in duration from control P_1_ cells, we conclude that the impact of losing PAR-1 is not due exclusively to its effect on cell size. Overall, our genetic and RNAi data are closely aligned and strongly suggest that PAR-1 activity is required for lower SAC activity in AB relative to P_1_.

## Discussion

We have shown that, during C. *elegans* embryogenesis, both cell size and cell fate impact the strength of the SAC. Spindle perturbations produce longer SAC-dependent mitotic delays in smaller cells, supporting the long-standing hypothesis that the strength of the SAC can be influenced by cell size (28, 29), while cells in the germline P lineage delay for longer than comparably-sized somatic cells, indicating that SAC strength is further subject to cell fate specific regulation. Variation in the strength of the SAC could arise at many points in the checkpoint signaling mechanism. Conceptually, we envision two primary possibilities: variation at the level of signal generation or variation in the rate of signal degradation. The former encompasses changes in the efficacy of MCC generation at the kinetochore or the number of MCC-generating kinetochores. The latter is a catchall class for changes that occur away from the kinetochore, including differences in the rate of cytoplasmic MCC disassembly or the efficiency of APC/C inhibition. A third possibility could arise downstream of the SAC, if, for example, cell size or cell fate changed the threshold of APC/C substrates at which cells exit mitosis. We think this unlikely, as the timing of mitotic exit is invariant across cells of different sizes and fates when SAC regulation is removed and APC/C activity is unimpeded.

Changes in cell size have been proposed to change the strength of the SAC via the first mechanism, increased signal generation. Smaller cells have a higher ratio of DNA to cytoplasm (28) and, effectively, a higher number of kinetochores per unit cytoplasm, thereby favoring MCC production (29). In cells with monopolar spindles, germline cell fate could act to increase the number of signal generating kinetochores. Cells with monopolar spindles fail to satisfy the SAC due to the persistence of monotelic and syntelic kinetochore-microtubule attachments (68). The presence of these attachments, albeit erroneous, may reduce the proportion of checkpoint-signaling kinetochores, as compared to Nocodazole treated cells, in which microtubule formation is largely blocked (26). Accordingly, longer monopolar mitoses in germline-fated cells could be achieved by further reducing kinetochore-microtubule attachment, thereby increasing the proportion of MCC-producing kinetochores. Alternatively, a microtubule-dependent checkpoint silencing mechanism functions at the kinetochore in *C. elegans* embryos (69) and rendering this mechanism more or less robust could also affect the proportion of checkpoint-signaling kinetochores in cells with monopolar spindles. While we cannot exclude either of these possibilities, we note that germline blastomeres may also exhibit longer mitotic arrests than somatic cells following Nocodazole treatment, in which very few kinetochore-microtubule attachments are made (29). Thus we favor the hypothesis that cell size sets the kinetochore to cytoplasmic ratio and cell fate acts in parallel, either at the level of signal generation, by modifying MCC production per kinetochore, or downstream of the kinetochore by modulating the cytoplasmic activity of the checkpoint.

Our results indicate that, at the 2-cell embryonic stage, PAR protein mediated cytoplasmic asymmetries underlie differential checkpoint strength between the somatic AB and germline P_1_ cell. PAR proteins regulate the levels and localization of certain asymmetrically distributed cellular components in 2-cell stage embryos largely via the posterior enrichment of PAR-1, which, in turn, drives anterior localization of two closely related proteins, MEX-5 and MEX-6 (hereafter MEX-5/6), and inhibits their enrichment and activity in the posterior (59). MEX-5/6 anchor and enrich certain factors in the anterior (66, 70) and promote the degradation of germline determinants in AB and/or its progeny (71). Removing anterior PAR proteins permits PAR-1 activity in the anterior, resulting in global inhibition of MEX-5/6 and two cells with certain germline P_1_ traits. Conversely, eliminating posterior PAR proteins allows MEX-5/6 activity in the posterior and certain somatic AB traits emerge in both cells (55).

Our results raise the question of why removing PAR-1, which is usually undetectable in AB, increased the duration of monopolar mitoses in AB, while depleting PAR-2 had a minimal effect. This result stands in contrast to a well-characterized asymmetric cellular behavior regulated by PAR proteins, cell cycle asynchrony (66, 70, 72, 73). If checkpoint activity were regulated in an analogous fashion, we would expect loss of *par-1* and *par-2* to result in shorter mitotic delays in P_1_, leaving AB largely unaffected. Instead, our results are consistent with a model in which a checkpoint-promoting factor (“factor X”) is enriched in P_1_ via a PAR-1-dependent mechanism. In the absence of PAR-2, anterior PAR proteins expand into the posterior, but are present in a graded fashion, and the posterior enrichment of some germline factors (e.g. P granules) is often maintained (58, 74). In addition, although PAR-1 is no longer asymmetric in *par-2(-)* embryos, its activity is not lost (58, 75, 76). As differential mitotic delays between AB and P_1_ cells in *par-2(lw32)* embryos were entirely dependent on PAR-1, we conclude that cytoplasmic asymmetries downstream of PAR-1, and partially independent of PAR-2, are largely responsible for posterior enrichment of factor X. In the absence of PAR-6, PKC-3 or PAR-1, this factor X is equally inherited by AB and P_1_, thereby increasing checkpoint activity in AB while maintaining it in P_1_ (Fig. 4e). This model requires that levels of factor X are not limiting for checkpoint activity and that factor X is not degraded in the absence of PAR-1, at least at the 2-cell stage. As in *par-1* mutants, degradation of germline factors is delayed such that germplasm proteins persist in all cells until the 4-cell stage (52), this latter condition is not without precedent.

Altogether, our results suggest that a checkpoint-regulating factor (or factors) is partitioned during the asymmetric division of germline blastomeres, downstream of PAR-mediated cell polarity, such that germline cells possess a stronger SAC relative to their nuclear to cytoplasmic ratio than their somatic siblings. SAC activity may be further tuned within each lineage as development progresses. Future work to identify the relevant molecular asymmetries between somatic and germline cells driving differential SAC activity will permit further investigation of this model and identification of where PAR-mediated cell polarity and checkpoint regulation intersect.

## Materials and Methods

### *C. elegans* strains and culture

All strains were maintained at 15°C on NGM plates, seeded with *E. coli* bacteria (OP50) following standard procedures (77). All genotypes are listed in Supplementary Table S1. L4 stage larvae were transferred to fresh OP50 plates at 15°C for 2 days, after which time embryos were harvested from gravid adults for imaging. RNAi depletions were performed at 15°C by feeding (78) as follows. Synchronized L1 larvae were obtained by sodium hypochlorite treatment (1.2% NaOCl, 250mM KOH) and were plated onto NGM plates containing 1.5mM IPTG and 25μg/ml Carbenicillin, seeded with HT115 bacteria containing the empty feeding vector L4440. Three days later early L4 larvae were transferred to fresh RNAi feeding plates seeded with bacteria expressing doubled-stranded RNA for gene inactivation or the empty vector L4440 for controls. Embryos were collected for imaging three days later. To isolate homozygous *par-2(lw32)* animals, L1 larvae on L4440 plates were shifted to 25°C for 24 hours, which allowed identification of homozygous animals based on the linked, recessive, temperature-sensitive allele *unc-45(e286)*. Unc larvae were transferred to RNAi feeding plates, returned to 15°C and imaged three days later. Control genotypes (UM471 or UM628, see Supplementary Table S1) were similarly treated. The following clones from the Arhinger RNAi library were used: sjj_C50F4.11 (*mdf-1*/*Mad1*), sjj_K10B2.5 (*ani-2*), sjj_ZK675.1 (*ptc-1*), sjj_C38C10.4 (*gpr-1/2*), sjj_T26E3.3 (*par-6*), sjj_F09E5.1 (*pkc-3*), sjj_H39E23.1 (*par-1*), and sjj_F58B6.3 (*par-2*). The clone targeting C27D9.1 was kindly provided by Dr. B. Lacroix. All clones were verified by sequencing.

### Embryo mounting and live-imaging

Embryos were harvested by cutting open gravid hermaphrodites in M9 buffer using two 25-gauge needles. Embryos were transferred by mouth pipet to a 2% agarose pad, positioned using an eyelash and covered with a coverslip. The chamber was backfilled with M9 and sealed using VaLaP (1:1:1 Vaseline, lanolin and paraffin). Embryos were mounted at room temperature in solutions that had been cooled to 15°C. Slides were transferred to a temperature-controlled imaging chamber set to 26°C and time-lapse images were acquired using either a Cell Observer SD spinning disc confocal (Zeiss; Yokogawa) equipped with a stage-top incubator (Pecon) or a DeltaVision inverted microscope (GE Healthcare Life Sciences) equipped with a WeatherStation environmental chamber (PrecisionControl). Spinning disc images were captured using a AxioCam 506 Mono camera (Zeiss) and a 63x/1.4 NA Plan Apochromat DIC oil immersion objective (Zeiss) in Zen software (Zeiss). DeltaVision images were captured using a Coolsnap HQ2 CCD camera (Photometrics) and a 60x/1.42 NA Plan Apo N oil immersion objective (Olympus) in softWoRx software (GE Healthcare Life Sciences). Time-lapse acquisitions were 45 to 60 minutes in duration, with approximately 30 second time sampling. At each time point a roughly 20μm thick z-stack was acquired, using either 1 or 1.5μm sectioning.

### Image processing and measurements

Image processing and analysis was carried out in ImageJ (NIH). The duration of bipolar, monopolar and *tbb-2(ts)* mitoses were determined manually by monitoring H2B::mCH fluorescence. NEBD was defined as the first frame in which non-incorporated H2B::mCH was lost from the nuclear area. DECOND was defined as the first frame in which the distribution of H2B::mCH shifted from bright and compact to fainter and diffuse. Maximal z-stack projections of unprocessed image files were scored. Single z-slices were examined if projected images were ambiguous. For all strains in which β-tubulin::GFP was also present (all experiments excepting *par-1(it51)* and its associated controls), the timing of NEBD was confirmed by the appearance of microtubules in the nuclear space and the start of DECOND was concomitant with the start of spindle disassembly and shrinking of the spindle pole. To exclude potentially confounding effects of accumulated mitotic errors due to any partial conditionality of our temperature sensitive alleles, only cells in which the preceding parental division was observed to be normal were analyzed. AB and P_1_ cell cycle duration was measured as the time from P_0_ metaphase to NEBD in each cell respectively, by manually monitoring H2B::mCH fluorescence. P_0_ metaphase was defined as the last time point prior to visible chromosome separation. NEBD was defined as above. To measure nuclear area, a sum projection of the central 3 z-slices of the nucleus of interest was processed and segmented. The average of the dimensions of a bounding box fit to the segmented nucleus was taken as representative of nuclear diameter and nuclear area was then calculated as the area of the corresponding circle (Supplementary Fig. S2a,b). Reported values reflect the average of measurements made at three time points, 1 to 2 minutes prior to NEBD. To determine the relative volume of cells in each lineage from the 2 to 16-cell stage, the position of the mitotic spindle midpoint along the division axis, relative to the midpoint of the dividing cell was used to determine what proportion of each dividing cell was allocated to each daughter (Supplementary Fig. S2c,d). The Plot Profile tool in ImageJ was used to generate signal intensity profiles along a line drawn parallel to the division axis, through the segregating sister nuclei, in cells in which the cell membrane was labeled with mNeonGreen::PH (mNG::PH) and nuclei were marked by H2B::mCH. Spindle displacement was defined as the offset of the center point between the two membrane mNG::PH peaks versus the center point between the two H2B::mCH peaks. When segregating sister nuclei were not in the same xy plane (i.e. the cell division axis was tilted relative to the plane of imaging), the Stack Reslice tool in ImageJ was used to construct an image of the long axis of the dividing cell from the encompassing z slices. Measurements were made at three time points during anaphase, after the start of membrane ingression. Nearly identical volume relationships were obtained when cell volume was calculated from cross-sectional area and cell height measurements using the mNG::PH membrane signal to manually outline cells (data not shown). The cell volume of AB and P_1_ was calculated by combining measurements of spindle displacement in P_0_, with measurements of embryo volume. Spindle displacement was measured using the Plot Profile tool in ImageJ to generate signal intensity profiles along the anterior to posterior axis of the embryo. The spindle midpoint was defined as the center point between the two peaks of β-tubulin::GFP fluorescence, corresponding to the spindle poles, and/or the two peaks of H2B::mCH fluorescence, corresponding to the segregating sister nuclei. The position of the spindle midpoint relative to the embryo midpoint was used as an indication of what proportion of the zygote would be inherited by each cell. Embryo volume was measured by manually outlining each embryo using the Polygon Selection Tool in ImageJ and fitting an ellipse to the resulting area. Embryo volume was then calculated as an ellipsoid of the same dimensions. Final values reflect the average of three measurements made at three different time points during anaphase in P_0_.

### Graphing and statistical analysis

Graphs were generated in Microsoft Excel or MATLAB (MathWorks). Statistical analysis was carried out in MATLAB. Comparison of multiple means was performed using an Anova1 with a Tukey-Kramer post-hoc test. Linear least squares regression models were calculated in MATLAB, and Pearson’s linear correlation coefficient (r) was reported, with statistical significance assessed using Student’s t-distribution. Regression coefficients were compared by bootstrapping, using a custom MATLAB script. For data presented in Fig. 3c,d, bootstrap analysis was performed manually in Excel, by calculating the difference in the duration of monopolar mitoses for all possible pairwise comparisons within a given size range. All possible outcomes were displayed as a boxplot graph and means were compared by an Anova1 with Tukey-Kramer post-hoc test. “Small” embryos are those in which AB is within ± 1 standard deviation (s.d.) of the size of a Control(RNAi) P_1_ cell. “Large” embryos are those in which P_1_ is within ± 1 s.d. of the size of a Control(RNAi) AB cell. “Normal” embryos are those in which AB is within ± 1 s.d. of the size of a Control(RNAi) AB cell. “P_1_” cell size compares AB and P_1_ cells that are both within ± 1 s.d. of the size of a Control(RNAi) P_1_ cell. “AB” cell size compares AB and P_1_ cells that are both within ± 1 s.d. of the size of a Control(RNAi) AB cell. For Fig. 3f,g, only *gpr-1/2*(RNAi), *par-6*(RNAi) and *par-1*(RNAi) embryos in which the position of the mitotic spindle in P_0_ was roughly centered (50±2% of embryo length) were considered. For data presented in Fig. 4b, for each RNAi treatment, AB and P_1_ cells were pooled and only cells within ±1 standard deviation of the average cell size for that depletion were considered. This same size range was applied to the total data set of normally polarized and fated AB and P_1_ cells (Control(RNAi), *ani-2*(RNAi), *ptc-1*(RNAi), *C27D9.1*(RNAi), *gpr-1/2*(RNAi)). Thus all cells, both depleted and control, AB and P_1_, analyzed for a given RNAi treatment are approximately the same size. For *par-1*(RNAi), the small size of the data set required increasing the size range to within ±2 standard deviations of the mean. For all Figures n.s. = p>0.05, * = p<0.05, ** = p<0.01 and *** = p<0.001.

### Author Contributions

A.R.G. designed, and carried out all experiments, with the exception of the *par-1(it51)* experiment which was carried out by V.P. A.R.G. analyzed the data, and wrote the manuscript with input from J-C.L. and P.S.M.

## Acknowledgements

We thank A. Golden, B. Goldstein, D. Dickinson and the *Caenorhabditis* Genetics Center for strains, C. Charbonneau of IRIC’s Bio-imaging facility for technical assistance and E. D. Salmon and members of the Labbé and Maddox labs for helpful discussions. This work was funded by a grant from the CIHR (MOP-115171) to J-C.L. and P.S.M. J-C.L. holds the Canada Research Chair in Cell Division and Differentiation. P.S.M. is supported as a William Burwell Harrison Fellow of Biology. IRIC is supported in part by the Canada Foundation for Innovation and the Fonds de Recherche du Québec – Santé.

## Supplemental Figures and Legends

**Figure S1.**
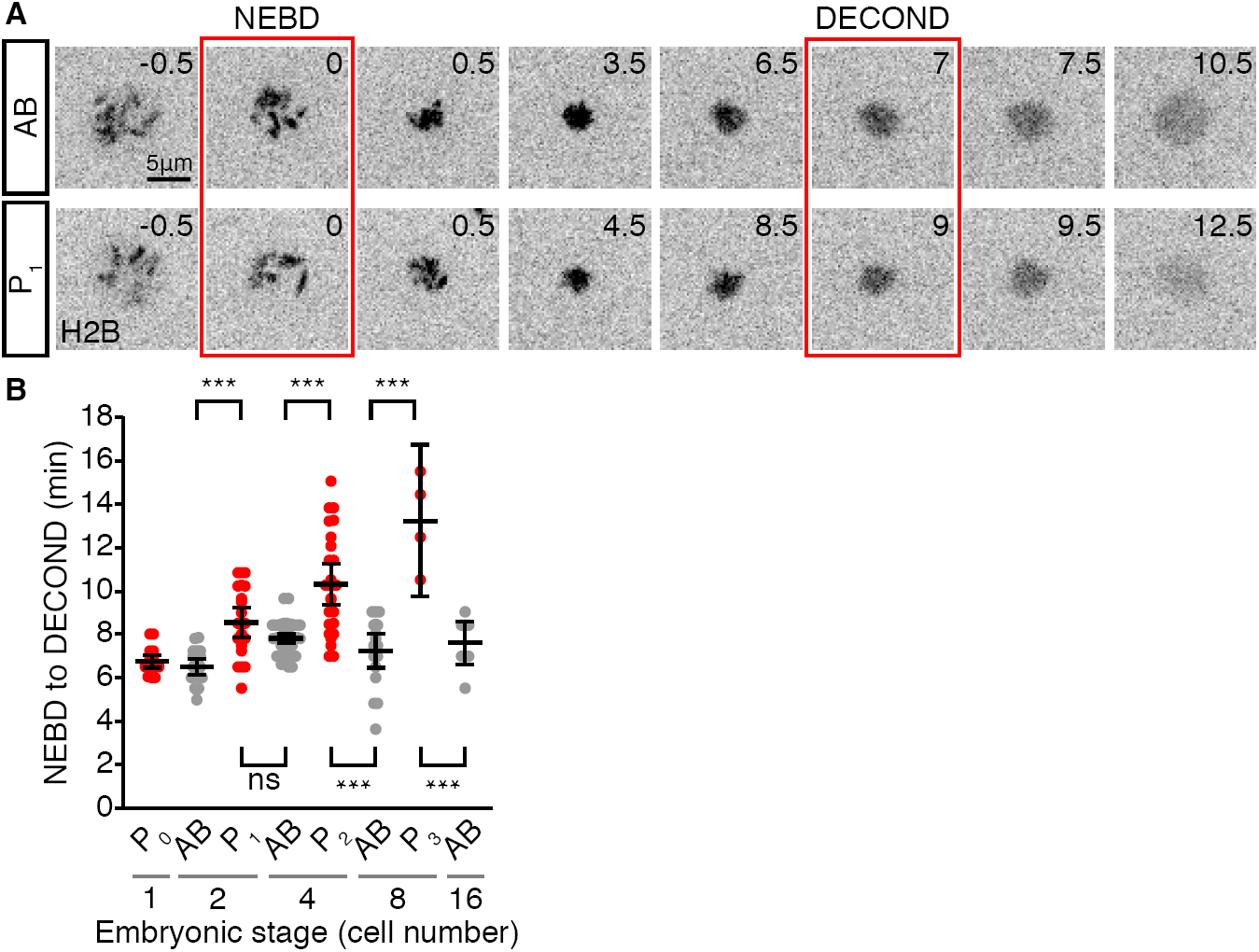
Germline P lineage cells show prolonged mitotic delays relative to somatic AB lineage cells when spindle formation is perturbed using a temperature sensitive allele of β-tubulin. (A) Single cropped time points from an AB (top) and P_1_ (bottom) cell carrying *tbb-2(or362)* and H2B::mCH and shifted to the restrictive temperature (26°C), showing the timing of nuclear envelop breakdown (NEBD), marked by the loss of non-chromatin associated H2B::mCH, to the start of chromosome decondensation (DECOND). (B) Graph showing the duration of mitosis in germline P and somatic AB lineage cells from 1 to 16-cell stage embryos with *tbb-2(or362)*-induced spindle perturbations. Germline P lineage cells (red) delay in mitosis for longer than somatic AB cells at the same and later embryonic stages. Black bars represent the mean. Error bars show the 95% confidence interval for the mean. p values were determined by an Anova1 with Tukey-Kramer post-hoc test, *** = p < 0.001 and ns = p > 0.05.

**Figure S2.**
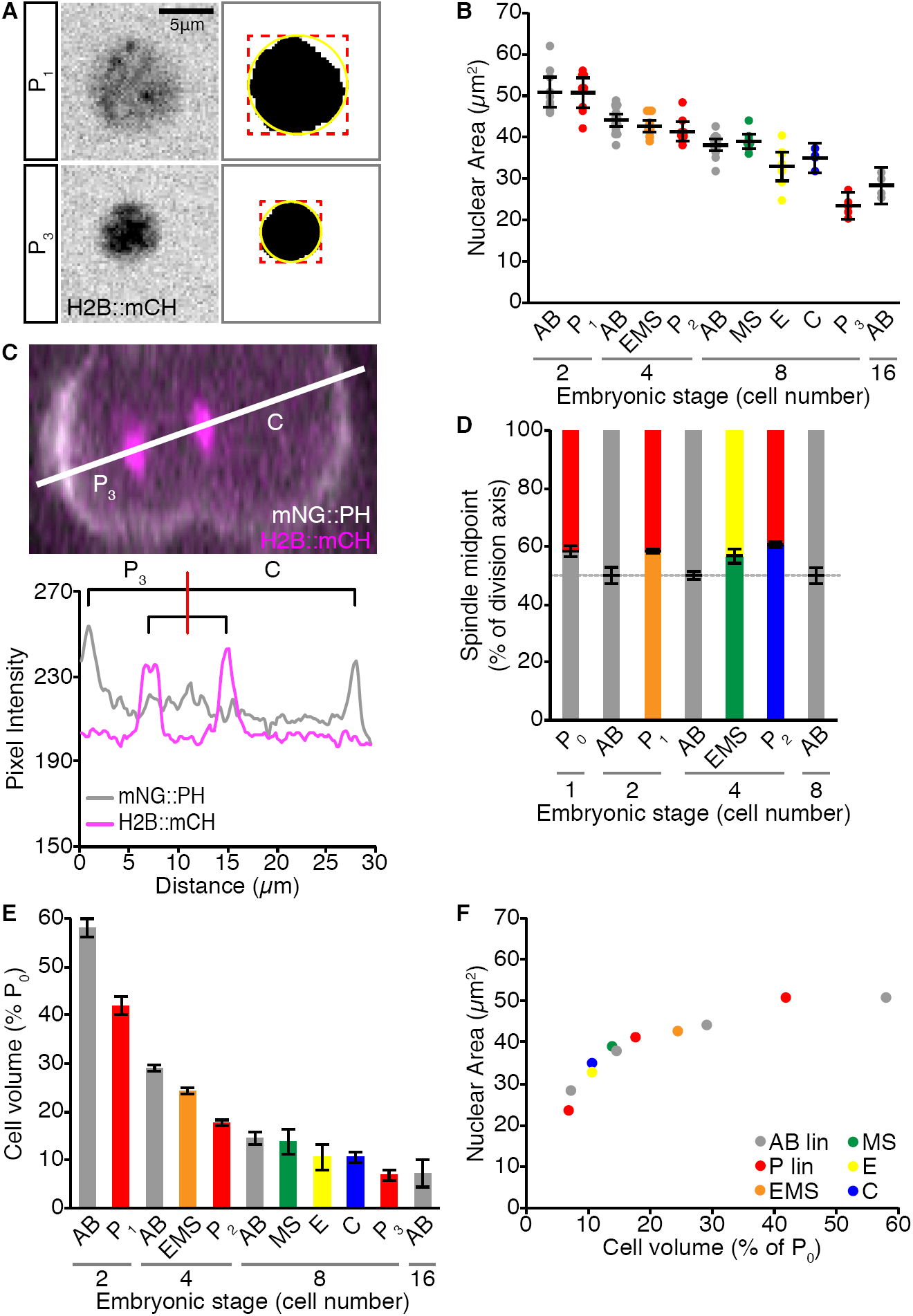
Nuclear area scales with cell volume as cell size decreases during embryonic development. (A) Sum projections through the center of an H2B::mCH-marked P_1_ (top) and P_3_ (bottom) nucleus one minute prior to NEBD, with the corresponding segmented image to the right. A bounding box was fit to the segmented nucleus and used to calculate the radius of a circle that approximates nuclear area. (B) Graph showing nuclear area measurements (performed as described in A) for the cells shown in Figures 1E and 2A. Black bars show the mean. Error bars represent the 95% confidence interval for the mean. (C) Image of a P_2_ cell division with the cell membrane marked by mNeonGreen::PH (mNG, white) and nuclei marked by H2B::mCH (magenta). White line shows the axis of cell division, along which spindle displacement was measured, with P_3_ to the left and C to the right. The pixel intensity values for each channel along this line are shown below. The spindle midpoint was measured as the center point between the two peaks of H2B::mCH fluorescence intensity and spindle displacement was calculated as the position of this center point relative to the center point of the long axis of the cell, as defined by the two fluorescence intensity peaks of the membrane marker mNG::PH. (D) Graph showing the symmetry or asymmetry of division, as measured by the method outlined in (C), for the different embryonic cells listed. Error bars show the standard deviation of the mean. (E) Graph showing cell volume for the different embryonic cells listed, represented as a percentage of the starting embryonic volume (P_0_) and calculated from the spindle displacement measurements shown in (D). Mean cell volume values were used in Figure 2B. (F) Graph showing the scaling relationship between mean nuclear area and mean cell volume.

**Figure S3.**
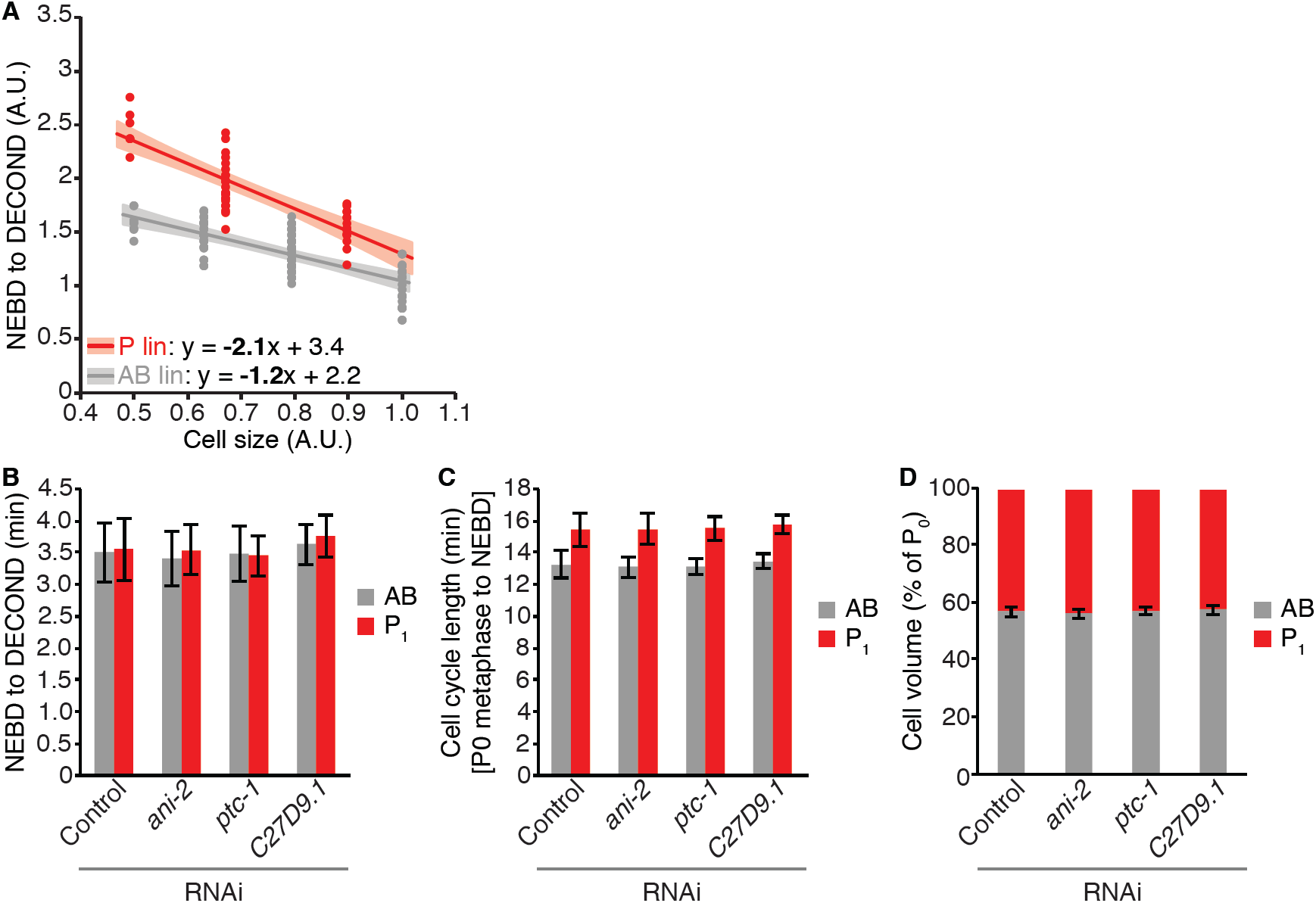
Cell fate and developmental stage impact the duration of monopolar mitoses. Scatter plot showing the duration of monopolar mitoses in cells from the germline P lineage (red) and somatic AB lineage (grey) in 2- to 16-cell embryos relative to mean cell size. Mean cell size was calculated by converting cell volume measurements from Figure S2E into the radius of the corresponding sphere (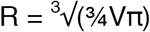, as in Figure 3B). Cells analyzed in (A) were treated with Control(RNAi) and are independent from those analyzed in Figure 2B. The rate at which the duration of monopolar mitoses increases with decreasing cell size is greater in the germline P lineage than in the somatic AB lineage (p = 0.02). (B-D) Bar graphs showing the mean duration of bipolar mitoses (B), cell cycle duration (C) and cell volume as a proportion of P_0_ (D) in AB (grey) and P_1_ (red) cells following RNAi-induced changes in embryo volume. Error bars represent ± 1 standard deviation of the mean.

**Figure S4.**
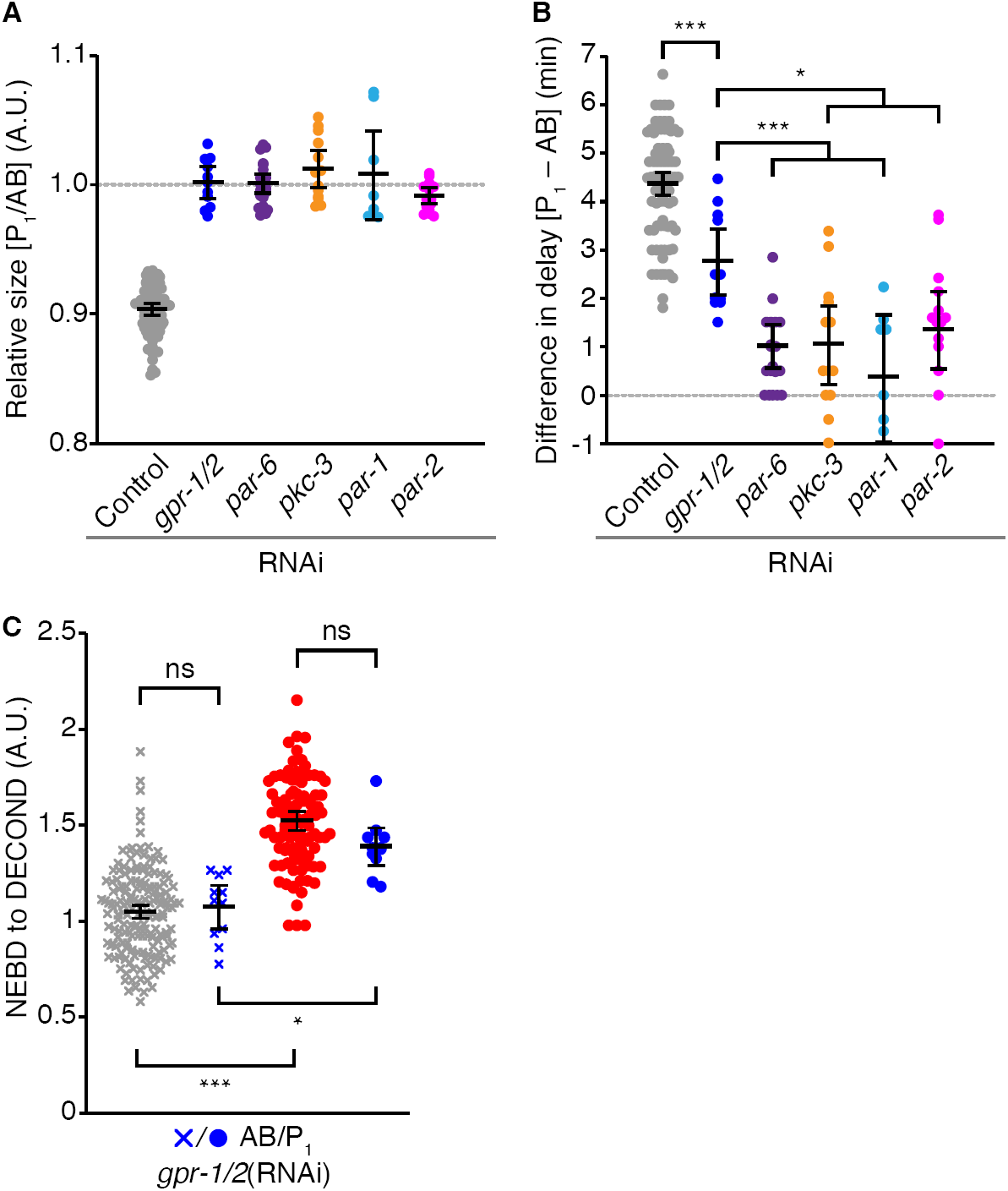
RNAi depletion of anterior and posterior PAR proteins reduces the difference in mitotic delay between AB/P1 sibling pairs more strongly than depletion of GPR-2. (A) Graph showing the size of the germline P_1_ cell relative to its somatic sibling, AB, following RNAi depletion of GPR-2 (blue), anterior (PAR-6 in purple and PKC-3 in orange) and posterior (PAR-1 in light blue and PAR-2 in magenta) PARs compared to Control RNAi (grey). (B) Graph showing the difference in the duration of monopolar mitoses between sibling AB and P_1_ cells for the same embryos shown in (A). Data for Control(RNAi), *gpr-2*(RNAi), *par-6*(RNAi) and *par-1*(RNAi) are reproduced from Figure 3F and 3G. (C) Graph comparing the duration of monopolar mitoses in somatic AB (Xs) and germline P_1_ (circles) cells following RNAi depletion of GPR-2, with comparably sized control cells. Control cells were pulled from data represented in Figure 3B. Both control and *gpr-2*(RNAi) cells were binned by size, with bin edges set at ± 2 standard deviations of the mean size for all *gpr-2*(RNAi) cells (both AB and P_1_). In both Control(RNAi) and *gpr-2*(RNAi) embryos, P_1_ cells have significantly longer monopolar mitoses than AB cells of the same size. *gpr-2*(RNAi) AB and *gpr-2*(RNAi) P_1_ cells have monopolar mitoses that are comparable in duration to similarly sized Control(RNAi) AB and P_1_ cells. For (A-C), black bars represent the mean and error bars show the 95% confidence interval. Data were compared using an Anova1 with Tukey-Kramer post-hoc test. *** = p < 0.001, * = p < 0.05, ns = p > 0.05.

**Figure S5.**
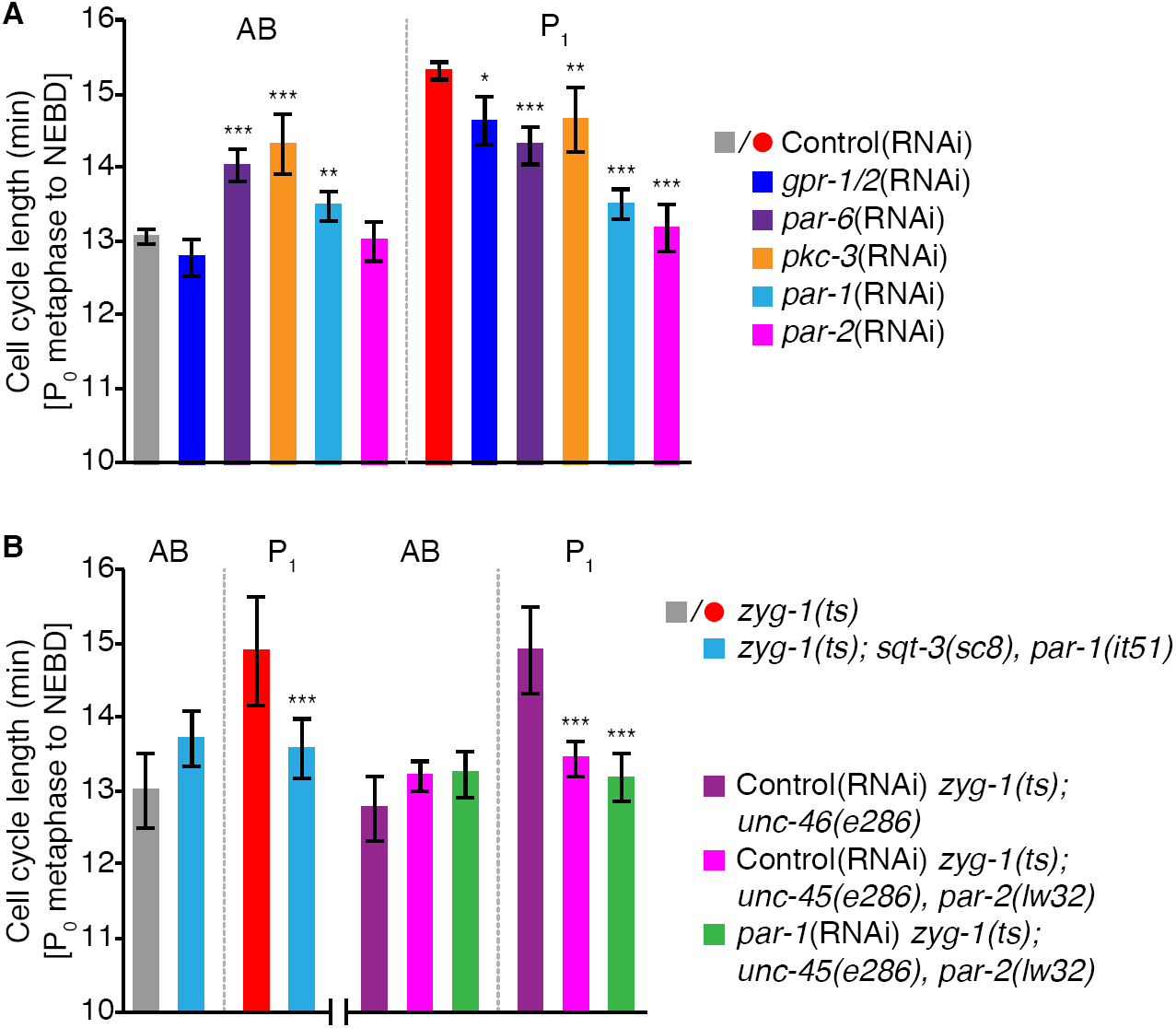
Cell cycle asynchrony is lost in *par* embryos, with depletion of anterior PAR and posterior PAR proteins delaying or accelerating cell cycle progression respectively. (A and B) Bar graphs showing the mean cell cycle length (from metaphase of P_0_ to nuclear envelope breakdown of AB or P_1_) for AB and P_1_. (A) Cell cycle length following RNAi depletion of GPR-1/2 (dark blue), PAR-6 (purple), PKC-3 (orange), PAR-1 (light blue) and PAR-2 (magenta). RNAi depletion of PAR-6 (purple) and PKC-3 (orange) increases cell cycle length in AB, while RNAi depletion of PAR-1 (light blue) and PAR-2 (magenta) decrease cell cycle length in P_1_. (B) Cell cycle lengths for *par-1(it51)* and *par-2(lw32)* AB and P_1_ cells, with their respective controls. Both *par-1(it51)* and *par-2(lw32)* P_1_ cells have shorter cell cycle durations than control P_1_ cells, while *par-1(it51)* and *par-2(lw32)* AB cells are unaffected. For (A and B), data were compared by an Anova1 with Tukey-Kramer post-hoc test and only statistically significant differences relative to control AB or P_1_ cells are shown. *** = p < 0.001, ** = p <0.01, and * = p < 0.05. Error bars show the 95% confidence interval for the mean.

## Supplemental Tables

**Table S1.**
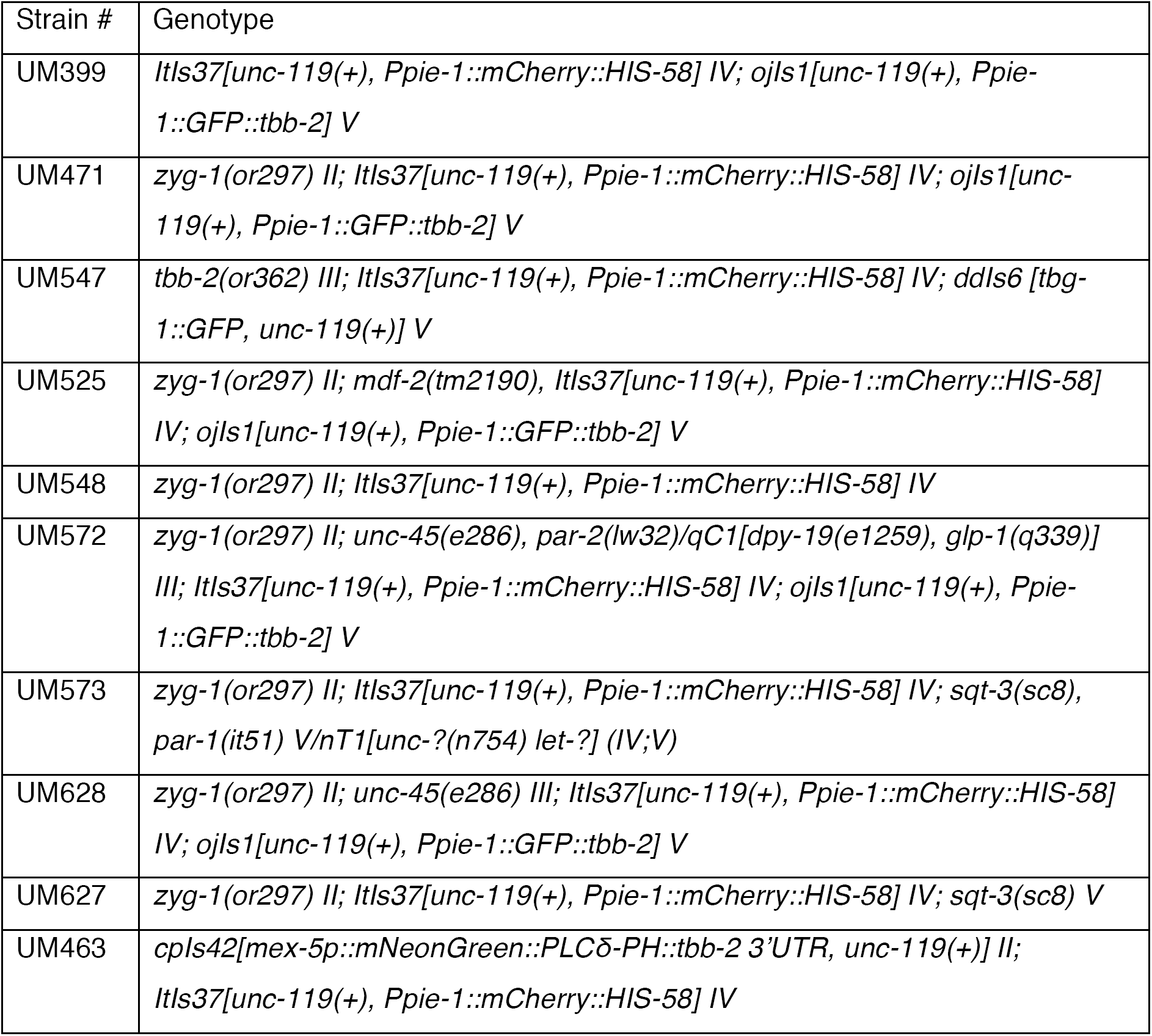
*C. elegans* strains used in this study. All strains, except UM399 (Gerhold 2015), were constructed as part of this study using the following: EU782 (*zyg-1(or297) II*), EU858 (*tbb-2(or362) III*), TH27 (*unc-119(ed3) III; ddIs6 [tbg-1::GFP, unc-119(+)] V*), UM397 (*mdf-2(tm2190), ItIs37[unc-119(+), Ppie-1::mCherry::his-58] IV*), LP271 (*cpIs42[mex-5p::mNeonGreen::PLCδ-PH::tbb-2 3’UTR + unc-119(+)]* II; unc-119(ed3) III), UM226 (*ItIs37[unc-119(+), Ppie-1::mCherry::his-58] IV*), KK747 (*unc-45(e286), par-2(lw32)/qC1[dpy-19(e1259), glp-1(q339)] III*), KK292 (*sqt-3(sc8), par-1(it51) V/nT1[unc-?(n754) let-?] (IV;V)*), BC8 (*sqt-3(sc8) V*), and CB286 (*unc-45(e286) III*). LP271 was kindly provided by Drs. D. Dickinson and B. Goldstein. The *mdf-2(tm2190)* allele was received from Dr. A. Golden. Some strains were provided by the CGC, which is funded by NIH Office of Research Infrastructure Programs (P40 OD010440).

## Supplemental Movies

All time-lapse movies were acquired on a Cell Observer SD spinning disc confocal (Zeiss; Yokogawa) equipped with a stage-top incubator (Pecon) set to 26°C, using an AxioCam 506 Mono camera (Zeiss), with 4x4 binning, and a 63x/1.4 NA Plan Apochromat DIC oil immersion objective (Zeiss) in Zen software (Zeiss). Images were acquired every 30 seconds, with the following exceptions: Movie S3 (*mdf-1*(RNAi)) every 33 seconds; Movie S4 (*gpr-1/2*(RNAi)), every 38 seconds. Images were processed using ImageJ (NIH). Maximum intensity projections of 18 z-slices (1 μm sectioning) centered on the embryo of interest are shown. Images were adjusted for brightness and contrast and scaled 4 times to adjust for file compression during .avi conversion. H2B::mCH is shown in cyan and β-tubulin::GFP is shown in red. All embryos are oriented with anterior to the left. Scale bar = 10 μm. Time stamp is in min:sec. Movies play at 4 frames per second, excepting Movie S4 (*gpr-1/2*(RNAi)), which plays at 3 frames per second.

**Movie S1. zyg-1(ts)-induced monopolar spindles in 2- and 4-cell stage embryos.**

**Movie S2. tbb-2(ts)-induced spindle defects in 2- and 4-cell stage embryos.**

**Movie S3. zyg-1(ts)-induced monopolar spindles in 2- and 4-cell stage embryos following RNAi depletion of MDF-1.**

**Movie S4. zyg-1(ts)-induced monopolar spindles in a 2-cell stage embryo following RNAi depletion of GPR-1/2.**

**Movie S5. zyg-1(ts)-induced monopolar spindles in a 2-cell stage embryo following RNAi depletion of PAR-6.**

## References

1. Musacchio A, Salmon ED. The spindle-assembly checkpoint in space and time. Nat Rev Mol Cell Biol. 2007;8(5):379–93.

2. Michel LS, Liberal V, Chatterjee A, Kirchwegger R, Pasche B, Gerald W, et al. MAD2 haplo-insufficiency causes premature anaphase and chromosome instability in mammalian cells. Nature. 2001;409(6818):355–9.

3. Cahill DP, Lengauer C, Yu J, Riggins GJ, Willson JK, Markowitz SD, et al. Mutations of mitotic checkpoint genes in human cancers. Nature. 1998;392(6673):300–3.

4. Weaver BA, Cleveland DW. Decoding the links between mitosis, cancer, and chemotherapy: The mitotic checkpoint, adaptation, and cell death. Cancer Cell. 2005;8(1):7–12.

5. Rieder CL, Maiato H. Stuck in division or passing through: what happens when cells cannot satisfy the spindle assembly checkpoint. Dev Cell. 2004;7(5):637–51.

6. Gascoigne KE, Taylor SS. Cancer cells display profound intra- and interline variation following prolonged exposure to antimitotic drugs. Cancer Cell. 2008;14(2):111–22.

7. Shi J, Orth JD, Mitchison T. Cell type variation in responses to antimitotic drugs that target microtubules and kinesin-5. Cancer Res. 2008;68(9):3269–76.

8. Clute P, Masui Y. Regulation of the appearance of division asynchrony and microtubule-dependent chromosome cycles in Xenopus laevis embryos. Dev Biol. 1995;171(2):273–85.

9. Clute P, Masui Y. Microtubule dependence of chromosome cycles in Xenopus laevis blastomeres under the influence of a DNA synthesis inhibitor, aphidicolin. Dev Biol. 1997;185(1):1–13.

10. Zhang M, Kothari P, Lampson MA. Spindle assembly checkpoint acquisition at the mid-blastula transition. PLoS One. 2015;10(3):e0119285.

11. Lara-Gonzalez P, Westhorpe FG, Taylor SS. The spindle assembly checkpoint. Curr Biol. 2012;22(22):R966–80.

12. London N, Biggins S. Signalling dynamics in the spindle checkpoint response. Nat Rev Mol Cell Biol. 2014;15(11):736–47.

13. Sudakin V, Chan GK, Yen TJ. Checkpoint inhibition of the APC/C in HeLa cells is mediated by a complex of BUBR1, BUB3, CDC20, and MAD2. J Cell Biol. 2001;154(5):925–36.

14. Chao WC, Kulkarni K, Zhang Z, Kong EH, Barford D. Structure of the mitotic checkpoint complex. Nature. 2012;484(7393):208–13.

15. Primorac I, Musacchio A. Panta rhei: the APC/C at steady state. J Cell Biol. 2013;201(2):177–89.

16. Clute P, Pines J. Temporal and spatial control of cyclin B1 destruction in metaphase. Nat Cell Biol. 1999;1(2):82–7.

17. Nasmyth K, Peters LM, Uhlmann F. Splitting the chromosome: Cutting the ties that bind sister chromatids. Science. 2000;288(5470):1379–84.

18. Mansfeld J, Collin P, Collins MO, Choudhary JS, Pines J. APC15 drives the turnover of MCC- CDC20 to make the spindle assembly checkpoint responsive to kinetochore attachment. Nat Cell Biol. 2011;13(10):1234–43.

19. Westhorpe FG, Tighe A, Lara-Gonzalez P, Taylor SS. p31comet-mediated extraction of Mad2 from the MCC promotes efficient mitotic exit. J Cell Sci. 2011;124(Pt 22):3905–16.

20. Lesage B, Qian J, Bollen M. Spindle checkpoint silencing: PP1 tips the balance. Curr Biol. 2011;21(21):R 898–903.

21. Howell BJ, McEwen BF, Canman JC, Hoffman DB, Farrar EM, Rieder CL, et al. Cytoplasmic dynein/dynactin drives kinetochore protein transport to the spindle poles and has a role in mitotic spindle checkpoint inactivation. J Cell Biol. 2001;155(7):1159–72.

22. Brito DA, Rieder CL. Mitotic checkpoint slippage in humans occurs via cyclin B destruction in the presence of an active checkpoint. Curr Biol. 2006;16(12):1194–200.

23. Joglekar AP. A Cell Biological Perspective on Past, Present and Future Investigations of the Spindle Assembly Checkpoint. Biology. 2016;5(4).

24. Musacchio A, Ciliberto A. The spindle-assembly checkpoint and the beauty of self-destruction. Nat Struct Mol Biol. 2012;19(11):1059–61.

25. Brito DA, Yang Z, Rieder CL. Microtubules do not promote mitotic slippage when the spindle assembly checkpoint cannot be satisfied. J Cell Biol. 2008;182(4):623–9.

26. Collin P, Nashchekina O, Walker R, Pines J. The spindle assembly checkpoint works like a rheostat rather than a toggle switch. Nat Cell Biol. 2013;15(11):1378–85.

27. Dick AE, Gerlich DW. Kinetic framework of spindle assembly checkpoint signalling. Nat Cell Biol. 2013;15(11):1370–7.

28. Minshull J, Sun H, Tonks NK, Murray AW. A MAP kinase-dependent spindle assembly checkpoint in Xenopus egg extracts. Cell. 1994;79(3):475–86.

29. Galli M, Morgan DO. Cell Size Determines the Strength of the Spindle Assembly Checkpoint during Embryonic Development. Dev Cell. 2016;36(3):344–52.

30. Gerhold AR, Ryan J, Vallee-Trudeau JN, Dorn JF, Labbe JC, Maddox PS. Investigating the regulation of stem and progenitor cell mitotic progression by in situ imaging. Curr Biol. 2015;25(9):1123–34.

31. Sulston JE, Schierenberg E, White JG, Thomson JN. The embryonic cell lineage of the nematode Caenorhabditis elegans. Dev Biol. 1983;100(1):64–119.

32. Bao Z, Zhao Z, Boyle TJ, Murray JI, Waterston RH. Control of cell cycle timing during C. elegans embryogenesis. Dev Biol. 2008;318(1):65–72.

33. Carvalho A, Olson SK, Gutierrez E, Zhang K, Noble LB, Zanin E, et al. Acute drug treatment in the early C. elegans embryo. PLoS One. 2011;6(9):e24656.

34. Strome S, Wood WB. Generation of asymmetry and segregation of germ-line granules in early C. elegans embryos. Cell. 1983;35(1):15–25.

35. O’Connell KF, Caron C, Kopish KR, Hurd DD, Kemphues KJ, Li Y, et al. The C. elegans zyg-1 gene encodes a regulator of centrosome duplication with distinct maternal and paternal roles in the embryo. Cell. 2001;105(4):547–58.

36. O’Rourke SM, Carter C, Carter L, Christensen SN, Jones MP, Nash B, et al. A survey of new temperature-sensitive, embryonic-lethal mutations in C. elegans: 24 alleles of thirteen genes. PLoS One. 2011;6(3):e16644.

37. Essex A, Dammermann A, Lewellyn L, Oegema K, Desai A. Systematic Analysis in Caenorhabditis elegans Reveals that the Spindle Checkpoint Is Composed of Two Largely Independent Branches. Mol Biol Cell. 2009;20(4):1252–67.

38. Arata Y, Takagi H, Sako Y, Sawa H. Power law relationship between cell cycle duration and cell volume in the early embryonic development of Caenorhabditis elegans. Front Physiol. 2014;5:529.

39. Ellis GC, Phillips JB, O’Rourke S, Lyczak R, Bowerman B. Maternally expressed and partially redundant beta-tubulins in Caenorhabditis elegans are autoregulated. J Cell Sci. 2004;117(Pt 3):457–64.

40. Edens LJ, White KH, Jevtic P, Li X, Levy DL. Nuclear size regulation: from single cells to development and disease. Trends Cell Biol. 2013;23(4):151–9.

41. Hara Y, Kimura A. Cell-size-dependent spindle elongation in the Caenorhabditis elegans early embryo. Curr Biol. 2009;19(18):1549–54.

42. Greenan G, Brangwynne CP, Jaensch S, Gharakhani J, Julicher F, Hyman AA. Centrosome size sets mitotic spindle length in Caenorhabditis elegans embryos. Curr Biol. 2010;20(4):353–8.

43. Ladouceur AM, Dorn JF, Maddox PS. Mitotic chromosome length scales in response to both cell and nuclear size. J Cell Biol. 2015;209(5):645–51.

44. Kitagawa R, Rose AM. Components of the spindle-assembly checkpoint are essential in Caenorhabditis elegans. Nat Cell Biol. 1999;1(8):514–21.

45. Consortium CeDM. large-scale screening for targeted knockouts in the Caenorhabditis elegans genome. G3. 2012;2(11):1415–25.

46. Tarailo-Graovac M, Wang J, Chu JS, Tu D, Baillie DL, Chen N. Spindle assembly checkpoint genes reveal distinct as well as overlapping expression that implicates MDF-2/Mad2 in postembryonic seam cell proliferation in Caenorhabditis elegans. BMC Cell Biol. 2010;11:71.

47. Green RA, Kao HL, Audhya A, Arur S, Mayers JR, Fridolfsson HN, et al. A high-resolution C. elegans essential gene network based on phenotypic profiling of a complex tissue. Cell. 2011;145(3):470–82.

48. Maddox AS, Habermann B, Desai A, Oegema K. Distinct roles for two C. elegans anillins in the gonad and early embryo. Development. 2005;132(12):2837–48.

49. Sonnichsen B, Koski LB, Walsh A, Marschall P, Neumann B, Brehm M, et al. Full-genome RNAi profiling of early embryogenesis in Caenorhabditis elegans. Nature. 2005;434(7032):462–9.

50. Guedes S, Priess JR. The C-elegans MEX-1 protein is present in germline blastomeres and is a P granule component. Development. 1997;124(3):731–9.

51. Jones AR, Francis R, Schedl T. GLD-1, a cytoplasmic protein essential for oocyte differentiation, shows stage- and sex-specific expression during Caenorhabditis elegans germline development. Dev Biol. 1996;180(1):165–83.

52. Reese KJ, Dunn MA, Waddle JA, Seydoux G. Asymmetric segregation of PIE-1 in C-elegans is mediated by two complementary mechanisms that act through separate PIE-1 protein domains. Mol Cell. 2000;6(2):445–55.

53. Schubert CM, Lin R, de Vries CJ, Plasterk RH, Priess JR. MEX-5 and MEX-6 function to establish soma/germline asymmetry in early C. elegans embryos. Mol Cell. 2000;5(4):671–82.

54. Seydoux G, Fire A. Soma-germline asymmetry in the distributions of embryonic RNAs in Caenorhabditis elegans. Development. 1994;120(10):2823–34.

55. Goldstein B, Macara IG. The PAR proteins: fundamental players in animal cell polarization. Dev Cell. 2007;13(5):609–22.

56. Guo S, Kemphues KJ. Par-1, a Gene Required for Establishing Polarity in C-Elegans Embryos, Encodes a Putative Ser/Thr Kinase That Is Asymmetrically Distributed. Cell. 1995;81(4):611–20.

57. Griffin EE. Cytoplasmic localization and asymmetric division in the early embryo of Caenorhabditis elegans. Wiley Interdiscip Rev Dev Biol. 2015;4(3):267–82.

58. Boyd L, Guo S, Levitan D, Stinchcomb DT, Kemphues KJ. PAR-2 is asymmetrically distributed and promotes association of P granules and PAR-1 with the cortex in C-elegans embryos. Development. 1996;122(10):3075–84.

59. Cuenca AA, Schetter A, Aceto D, Kemphues K, Seydoux G. Polarization of the C. elegans zygote proceeds via distinct establishment and maintenance phases. Development. 2003;130(7):1255–65.

60. Etemad-Moghadam B, Guo S, Kemphues KJ. Asymmetrically distributed PAR-3 protein contributes to cell polarity and spindle alignment in early C. elegans embryos. Cell. 1995;83(5):743–52.

61. Griffin EE, Odde DJ, Seydoux G. Regulation of the MEX-5 gradient by a spatially segregated kinase/phosphatase cycle. Cell. 2011;146(6):955–68.

62. Kemphues KJ, Priess JR, Morton DG, Cheng NS. Identification of genes required for cytoplasmic localization in early C. elegans embryos. Cell. 1988;52(3):311–20.

63. Colombo K, Grill SW, Kimple RJ, Willard FS, Siderovski DP, Gonczy P. Translation of polarity cues into asymmetric spindle positioning in Caenorhabditis elegans embryos. Science. 2003;300(5627):1957–61.

64. Gotta M, Dong Y, Peterson YK, Lanier SM, Ahringer J. Asymmetrically distributed C. elegans homologs of AGS3/PINS control spindle position in the early embryo. Curr Biol. 2003;13(12):1029–37.

65. Srinivasan DG, Fisk RM, Xu H, van den Heuvel S. A complex of LIN-5 and GPR proteins regulates G protein signaling and spindle function in C elegans. Genes Dev. 2003;17(10):1225–39.

66. Rivers DM, Moreno S, Abraham M, Ahringer J. PAR proteins direct asymmetry of the cell cycle regulators Polo-like kinase and Cdc25. J Cell Biol. 2008;180(5):877–85.

67. Levitan DJ, Boyd L, Mello CC, Kemphues KJ, Stinchcomb DT. par-2, a gene required for blastomere asymmetry in Caenorhabditis elegans, encodes zinc-finger and ATP-binding motifs. Proc Natl Acad Sci U S A. 1994;91(13):6108–12.

68. Kapoor TM, Mayer TU, Coughlin ML, Mitchison TJ. Probing spindle assembly mechanisms with monastrol, a small molecule inhibitor of the mitotic kinesin, Eg5. J Cell Biol. 2000;150(5):975–88.

69. Espeut J, Cheerambathur DK, Krenning L, Oegema K, Desai A. Microtubule binding by KNL-1 contributes to spindle checkpoint silencing at the kinetochore. J Cell Biol. 2012;196(4):469–82.

70. Nishi Y, Rogers E, Robertson SM, Lin R. Polo kinases regulate C. elegans embryonic polarity via binding to DYRK2-primed MEX-5 and MEX-6. Development. 2008;135(4):687–97.

71. DeRenzo C, Reese KJ, Seydoux G. Exclusion of germ plasm proteins from somatic lineages by cullin-dependent degradation. Nature. 2003;424(6949):685–9.

72. Noatynska A, Tavernier N, Gotta M, Pintard L. Coordinating cell polarity and cell cycle progression: what can we learn from flies and worms? Open Biol. 2013;3(8):130083.

73. Budirahardja Y, Gonczy P. PLK-1 asymmetry contributes to asynchronous cell division of C. elegans embryos. Development. 2008;135(7):1303–13.

74. Beatty A, Morton DG, Kemphues K. PAR-2, LGL-1 and the CDC-42 GAP CHIN-1 act in distinct pathways to maintain polarity in the C. elegans embryo. Development. 2013;140(9):2005–14.

75. Labbe JC, Pacquelet A, Marty T, Gotta M. A genomewide screen for suppressors of par-2 uncovers potential regulators of PAR protein-dependent cell polarity in Caenorhabditis elegans. Genetics. 2006;174(1):285–95.

76. Watts JL, Etemad-Moghadam B, Guo S, Boyd L, Draper BW, Mello CC, et al. par-6, a gene involved in the establishment of asymmetry in early C. elegans embryos, mediates the asymmetric localization of PAR-3. Development. 1996;122(10):3133–40.

77. Brenner S. The genetics of Caenorhabditis elegans. Genetics. 1974;77(1):71–94.

78. Kamath RS, Martinez-Campos M, Zipperlen P, Fraser AG, Ahringer J. Effectiveness of specific RNA-mediated interference through ingested double-stranded RNA in Caenorhabditis elegans. Genome Biol. 2001;2(1):Research0002.

